# Sustained bacterial N_2_O reduction at acidic pH

**DOI:** 10.1101/2023.10.06.560748

**Authors:** Guang He, Gao Chen, Yongchao Xie, Cynthia Swift, Gyuhyon Cha, Konstantinos T. Konstantinidis, Mark Radosevich, Frank E. Löffler

**Author notes:** Frank E. Löffler, University of Tennessee, Department of Civil and Environmental Engineering, 325 John D. Tickle Building, 851 Neyland Drive, Knoxville, TN 37996, USA. **Email:**. Department of Chemistry and Biochemistry, University of California, Los Angeles, Los Angeles, CA 90095, USA.

## Abstract

Nitrous oxide (N_2_O) is a climate-active gas and emissions from terrestrial ecosystems are concerning. Microbial reduction of N_2_O to dinitrogen (N_2_) is the only known consumption process and has been studied extensively at circumneutral pH; however, N_2_O reduction under acidic conditions is thought to be limited. Global soil acidification, accelerated by anthropogenic practices, introduces high uncertainty into N_2_O emission budgets. We obtained an enrichment culture from an acidic tropical forest soil that robustly reduces N_2_O to N_2_ at pH 4.5 with the addition of pyruvate and hydrogen. Consecutive transfers at pH 4.5 yielded a co-culture and temporal analyses revealed a bimodal growth pattern with a *Serratia* sp. growing during the initial pyruvate fermentation phase followed by growth of a novel *Desulfosporosinus* sp. via hydrogenotrophic N_2_O reduction. The *Desulfosporosinus* sp. produced (3.1 ± 0.11) × 10^8^ cells per mmol of N_2_O consumed, on par with growth yields reported for clade II N_2_O reducers at circumneutral pH. Genome analysis identified a clade II *nos* gene cluster, but an incomplete pathway for sulfate reduction, a hallmark feature of the genus *Desulfosporosinus*. Physiological and metabogenomic characterization revealed interspecies nutritional interactions, with the pyruvate fermenting *Serratia* sp. supplying amino acids as essential growth factors to the *Desulfosporosinus* sp. The co-culture reduced N_2_O between pH 4.5 and 6 but not at or above pH 7, contradicting the paradigm that sustained microbial N_2_O reduction ceases under acidic pH conditions, thus confirming a previously unrecognized N_2_O reduction potential in acidic soils.

**Significance Statement:** Processes generating N_2_O occur over a broad pH range spanning pH 3 to 12; however, the current paradigm assumes that microbial N_2_O consumption is limited to circumneutral pH (6 to 8). The imbalance between N_2_O production versus consumption has increased the atmospheric concentration of this climate active gas by 17 % over the last 100 years, and accelerated emissions due to global soil acidification are a major climate concern. From acidic soil, we obtained a bacterial culture harboring a novel *Desulfosporosinus* species that effectively reduces N_2_O at pH 4.5, but not at or above pH 7. The discovery of an N_2_O reducer adapted to acidic pH conditions has far-reaching implications for predicting, modeling, and potentially managing N_2_O emissions from low pH ecosystems.

**Note for publisher (this text will be removed prior to publication):** This manuscript has been authored by UT-Battelle, LLC under Contract No. DE-AC05-00OR22725 with the U.S. Department of Energy. The United States Government retains and the publisher, by accepting the article for publication, acknowledges that the United States Government retains a non-exclusive, paid-up, irrevocable, world-wide license to publish or reproduce the published form of this manuscript, or allow others to do so, for United States Government purposes. The Department of Energy will provide public access to these results of federally sponsored research in accordance with the DOE Public Access Plan (http://energy.gov/downloads/doe-public-access-plan).

## Introduction

Exploitation, mismanagement, and unsustainable practices deteriorate soil health across the globe (1). pH is a key soil health parameter, but soil acidification, a natural process accelerated by the reliance of synthetic nitrogen fertilizer, the growth of legumes, and acidic precipitation/deposition, plagues regions around the world (2, 3). In addition to pH change, N fertilizer input substantially increases the formation of N_2_O, a compound linked to ozone depletion and climate warming (4–8), and interference with biogeochemical processes such as methanogenesis, mercury methylation, and reductive dechlorination (9, 10). The rise in global N_2_O emissions indicates an imbalance between N_2_O formation versus consumption, which has been explained by the contrasting pH optima of processes involved in N_2_O formation (pH 5-6) versus consumption (circumneutral pH) (11). A few studies reported N_2_O consumption in acidic soil laboratory microcosms (12, 13); however, soil heterogeneity and associated (microscale) patchiness of pH conditions make generalized conclusions untenable (14, 15). Attempts with denitrifying enrichment and axenic cultures have thus far failed to demonstrate growth-linked N_2_O reduction and sustainability of such a process under acidic pH conditions (15–17).

The only known sink for N_2_O are microorganisms expressing N_2_O reductase (NosZ), a periplasmic enzyme that catalyzes the conversion of N_2_O to environmentally benign dinitrogen (N_2_). NosZ expression and proteomics studies with the model denitrifier *Paracoccus denitrificans* demonstrated that acidic pH interferes with NosZ maturation (18, 19), a phenomenon also observed in enrichment cultures harboring diverse N_2_O-reducing bacteria (20). Studies with *Marinobacter hydrocarbonoclasticus* found active NosZ with a Cu_Z_ center in the 4Cu2S form in cells grown at pH 7.5, but observed a catalytically inactive NosZ with the Cu_Z_ center in the form 4Cu1S when the bacterium was grown at pH 6.5 (21). These observations support the paradigm that microbial N_2_O reduction is limited to circumneutral pH environments. A metagenome-based analysis of soil microbial communities in the Luquillo Experimental Forest (Puerto Rico) provided evidence that N_2_O-reducing soil microorganisms are not limited to circumneutral pH soils and exist in acidic tropical forest soils (22). Microcosms established with acidic Luquillo Experimental Forest soil demonstrated the N_2_O reduction activity, and suggested that the microbial communities in pH 4.5 and pH 7 microcosms differed (23). To shed light on the microorganisms performing N_2_O reduction at pH 4.5, a two-population co-culture was derived from N_2_O-reducing soil microcosms established with acidic tropical forest soil collected in the Luquillo Experimental Forest (El Yunque National Forest) in Puerto Rico. Characterization of the co-culture demonstrated the activity of an acidophilic *Desulfosporosinus* sp. that utilizes N_2_O as electron acceptor at pH 4.5.

## Results

### A consortium consisting of two populations reduces N_2_O at pH 4.5

Microcosms established with El Verde tropical soil amended with lactate consumed N_2_O at pH 4.5; however, this activity was lost in transfer cultures amended with the same electron donor. The addition of acetate, formate (1 or 5 mM each), and CO_2_ (208 µmol, 2.08 mM nominal), propionate (5 mM), or yeast extract (0.10 – 10 g L^-1^) did not stimulate N_2_O reduction in pH 4.5 transfer cultures. Limited N_2_O consumption was observed in transfer cultures amended with 2.5 mM pyruvate, but complete removal of N_2_O required the addition of H_2_ or formate. In transfer cultures with H_2_ or formate, but lacking pyruvate, N_2_O was not consumed. Subsequent transfers in completely synthetic basal salt medium amended with both pyruvate and H_2_ yielded a robust enrichment culture, which consumes N_2_O at pH 4.5 (Fig. 1A). Phenotypic characterization illustrated that pyruvate utilization was independent of N_2_O, while N_2_O reduction only commenced following complete consumption of pyruvate. The fermentation of pyruvate in culture EV yielded acetate, CO_2_, and formate as measurable products, with formate and external H_2_ serving as electron donors for subsequent N_2_O reduction (*SI Appendix*, Fig. S1 and Text S1). The fermentation of pyruvate resulted in pH increases, with the magnitude of the medium pH change proportional to the initial pyruvate concentration. The amount of pyruvate did not exceed 250 µmol per 100 mL of medium (i.e., 2.5 mM), which limited the pH increase following complete pyruvate fermentation to 0.53 ± 0.03 pH units (*SI Appendix*, Fig. S2). N_2_O reduction was also observed in cultures that received 5 mM glucose. N_2_O reduction was oxygen sensitive and N_2_O was not consumed in medium without reductant (i.e., cysteine, dithiothreitol).

**Fig. 1.**
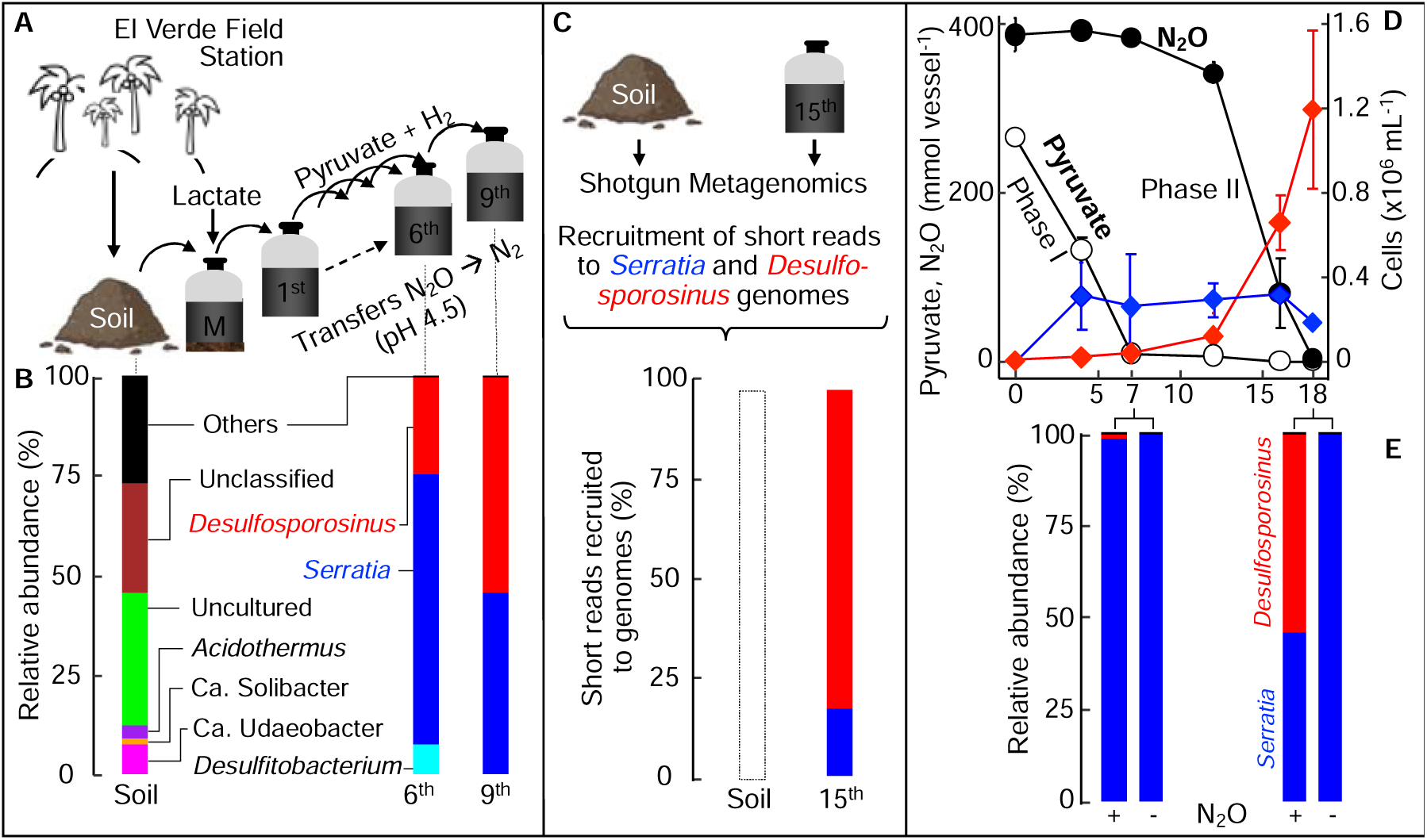
Establishment of N_2_O-reducing microcosms and enrichment cultures yielding co-culture EV. (A) Schematic of the workflow leading from a soil sample to a solids-free enrichment, and to a co-culture. (B) Community structure profiling based on 16S rRNA gene sequence analysis documents the enrichment process. Soil microbial community structure was profiled with 16S rRNA genes extracted from shotgun metagenomic reads. Community profiling of 6^th^ and 9^th^ transfers was based on 16S rRNA gene amplicon sequencing. Sequences with abundances <2% were grouped as “Others”. (C) Percent of the metagenomic short read fragments obtained from El Verde soil and the 15^th^ transfer culture that recruited to the genomes of *Serratia* sp. or *Desulfosporosinus* sp. A representative graph showing the high identity (> 95%) of reads mapping evenly across *Desulfosporosinus* sp. genome is presented in Fig. S3. The *Serratia* sp. or *Desulfosporosinus* sp. genomes were not detected in the soil metagenome dataset, with less than 0.01% of the metagenomic reads mapping to the two genomes (white bar). (D) Pyruvate fermentation (Phase I, white circles) and N_2_O consumption (Phase II, black circles) in co-culture EV. *Serratia* (blue diamonds) and *Desulfosporosinus* (red diamonds) cell numbers were determined with specific, 16S rRNA gene-targeted qPCR. (E) Amplicon sequencing illustrates the population shifts in co-culture EV following pyruvate consumption (day 7) and following N_2_O consumption (day 18). Relative abundance of *Serratia* (blue bars) and *Desulfosporosinus* (red bars) in co-culture EV following Phase I (day 7) and Phase II (day 18) in vessels with pyruvate plus N_2_O (left) and pyruvate only (right). Sequencing was performed on single representative cultures. All other data represent the averages of triplicate incubations and error bars represent standard deviations (n=3). Error bars are not shown if smaller than the symbol.

Microbial profiling of El Verde soil and solids-free transfer cultures documented effective enrichment in defined pH 4.5 medium amended with pyruvate, H_2_, and N_2_O (Fig. 1 B and *SI Appendix*, Text S2). Following nine consecutive transfers, *Serratia* and *Desulfosporosinus* each contributed about half of the 16S rRNA amplicon sequences (49.7% and 50.2%, respectively), and less than 0.05% of the sequences represented *Planctomycetota*, *Lachnoclostridium*, *Caproiciproducens*.

Deep shotgun metagenome sequencing performed on a 15^th^ transfer culture recovered two draft genomes representing the *Serratia* sp. and the *Desulfosporosinus* sp., accounting for more than 95% of the total short read fragments. All 16S rRNA genes associated with assembled contigs could be assigned to *Serratia* or *Desulfosporosinus* (*SI Appendix*, Fig. S4, Text S2), indicating that the enrichment process yielded a consortium consisting of a *Serratia* sp. and a *Desulfosporosinus* sp., designated co-culture EV. Efforts to recover the *Serratia* and *Desulfosporosinus* genomes from the original soil metagenome data sets via recruiting the soil metagenome fragments to the two genomes (Fig. 1 C) were not successful, highlighting the effectiveness of the enrichment strategy. Redundancy-based analysis with Nonpareil (24) revealed that the average covered species richness in the metagenome data set obtained from the 15^th^ transfer culture was 99.9%, much higher than what was achieved for the El Verde soil (39.5%), suggesting the metagenome analysis of the original soil did not fully capture the resident microbial diversity.

The application of 16S rRNA gene-targeted qPCR assays to DNA extracted from 9^th^ transfer N_2_O-reducing cultures revealed a bimodal growth pattern. During pyruvate fermentation (Phase I), the *Serratia* cell numbers increased nearly 1,000-fold from (2.3 ± 0.8) × 10^2^ to (1.8 ± 0.2) × 10^5^ cells mL^-1^, followed by a 40-fold increase from (3.5 ± 1.5) × 10^4^ to (1.2 ± 0.4) × 10^6^ cells mL^-1^ of *Desulfosporosinus* cells during N_2_O reduction (Phase II) (Fig. 1 D). In vessels without N_2_O, *Desulfosporosinus* cell numbers did not increase, indicating that growth of this population depended on the presence of N_2_O. Growth yields of (3.1 ± 0.11) × 10^8^ cells mmol^-1^ of N_2_O and (7.0 ± 0.72) × 10^7^ cells mmol^-1^ of pyruvate were determined for the *Desulfosporosinus* and the *Serratia* populations, respectively. 16S rRNA gene amplicon sequencing performed on representative samples collected at the end of Phase I (day 7) and Phase II (day 18) confirmed a bimodal growth pattern. Sequences representing *Serratia* increased during Phase I and *Desulfosporosinus* sequences increased during Phase II (Fig. 1 E). Taken together, the physiological characterization, qPCR, genomic, and amplicon sequencing results indicate that co-culture EV performs low pH N_2_O reduction, with a *Serratia* sp. fermenting pyruvate and a *Desulfosporosinus* sp. reducing N_2_O. Isolation efforts yielded an axenic *Serratia* sp., designated strain MF, capable of pyruvate fermentation. Despite extensive efforts, the *Desulfosporosinus* sp. resisted isolation, presumably due to auxotrophic interactions with strain MF (*SI Appendix*, Fig. S5 and Text S3).

Identification of auxotrophies.

To investigate the specific nutritional requirements of the *Desulfosporosinus* sp. in co-culture EV, untargeted metabolome analysis was conducted on supernatant collected from axenic *Serratia* sp. cultures growing with pyruvate and during N_2_O consumption (Phase II) following inoculation with co-culture EV (Fig. 2 A). Peaks representing potential metabolites were searched against a custom library (*SI Appendix*, Table S5) and 33 features were assigned to known structures, including seven amino acids (alanine, glutamate, methionine, valine, leucine, aspartate, and tyrosine). Cystine, the oxidized derivative of the amino acid cysteine, was also detected; however, cystine or cysteine were not found in cultures where dithiothreitol (DTT) replaced cysteine as the reductant, suggesting that *Serratia* did not excrete either compound into the culture supernatant. Time series metabolome analysis of culture supernatant demonstrated dynamic changes to the amino acid profile following inoculation with the *Serratia* sp. and the *Desulfosporosinus* sp. (as co-culture EV) (Fig. 2 A, B). Alanine, valine, leucine, and aspartate increased during pyruvate fermentation (Phase I) and were not consumed by the *Serratia* sp. (*SI Appendix*, Fig. S6). Consumption of alanine, valine, leucine, and aspartate did occur following the inoculation of the *Desulfosporosinus* sp. (as co-culture EV) (Fig. 2 A).

**Fig. 2.**
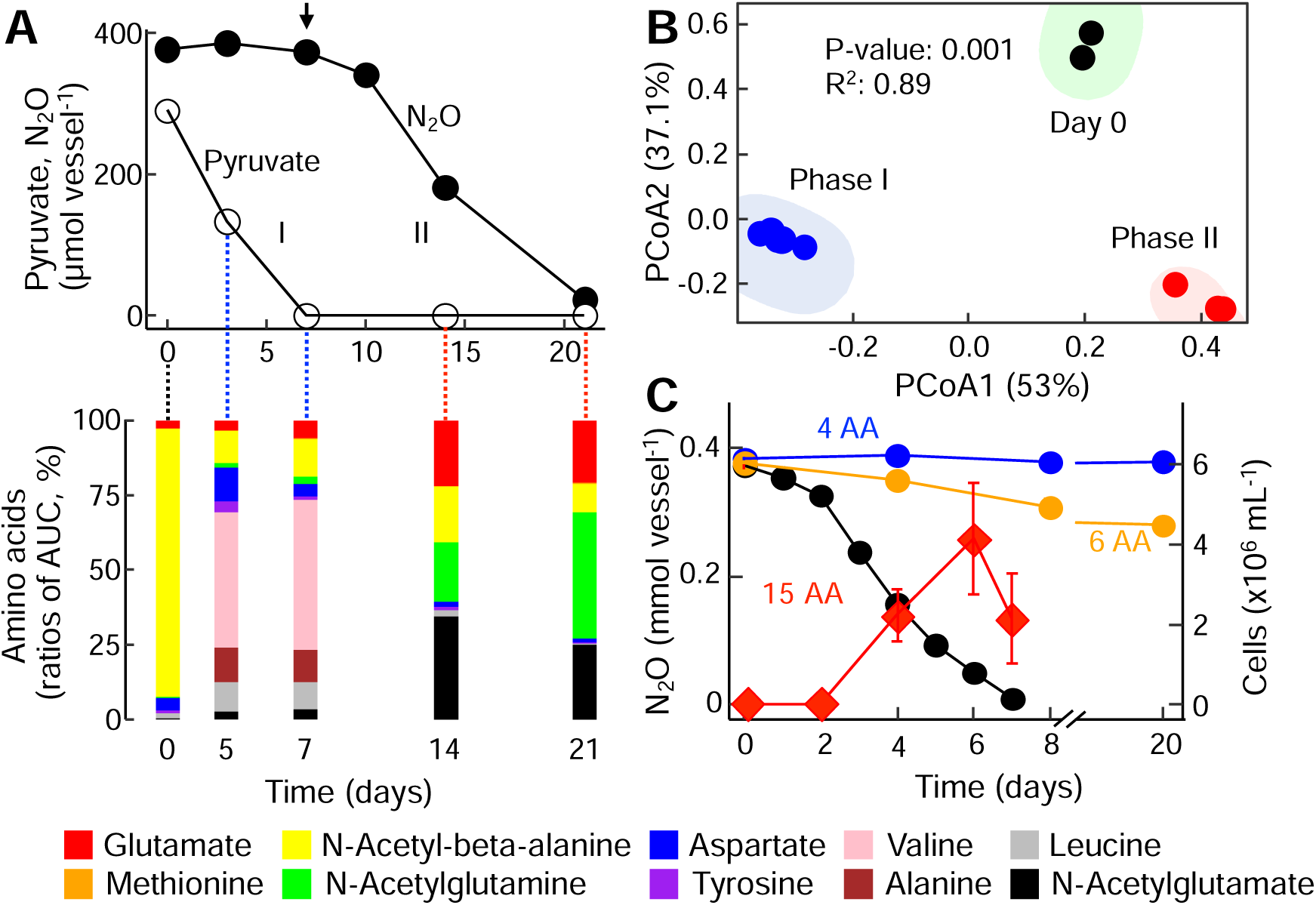
Interspecies cross-feeding supports low pH N_2_O reduction in co-culture EV. (A) Pyruvate fermentation (Phase I, light blue background) in vessels inoculated with axenic *Serratia* sp. strain MF and N_2_O consumption (Phase II, red background) following inoculation with co-culture EV comprising the *Serratia* sp. and the N_2_O-reducing *Desulfosporosinus* sp. (indicated by the arrow). The bottom part of Panel A shows the amino acid profile in the supernatant immediately after inoculation with the *Serratia* sp. (green background), during Phase I, and during Phase II following inoculation with co-culture EV (day 7; 3% inoculum). Samples for untargeted metabolome analysis were collected immediately after inoculation with *Serratia* sp. (light green), during Phase I (light blue, pyruvate fermentation) and Phase II (light red, N_2_O reduction). The stacked bars show the ratio (%) of areas under the curve (AUC) of the respective amino acids and amino acid derivatives. Metabolites not assigned to structures representing amino acids or its derivatives are not shown. (B) Principal coordinate analysis (PCoA) of amino acid profiles. The enclosing ellipses were estimated using the Khachiyan algorithm with the ggforce package. Permanova analysis was conducted with the vegan Community Ecology package. Black, blue, and red circles represent samples collected at day 0, during Phase I, and during Phase II, respectively. (C) N_2_O consumption in co-culture EV in medium amended with mixtures comprising 5 (blue), 6 (orange), or 15 (black) amino acids (see SI for composition of mixtures), H_2_, and N_2_O. The *Desulfosporosinus* sp. cell numbers (red diamonds) were determined with 16S rRNA gene-targeted qPCR and show growth in medium receiving the 15-amino acid mixture. Various other amino acid mixtures tested resulted in no or negligible N_2_O consumption. All growth assays with amino acid mixture supplementation were performed in triplicates and repeated in independent experiments.

These findings suggest that the N_2_O-reducing *Desulfosporosinus* sp. is an amino acid auxotroph, and a series of growth experiments explored if amino acid supplementation (*SI Appendix*, Table S6) can replace the requirement for pyruvate fermentation by the *Serratia* sp. for enabling subsequent N_2_O consumption by the *Desulfosporosinus* sp. The addition of individual amino acids (n=20) was not sufficient to initiate N_2_O reduction in pH 4.5 medium, as was the combination of alanine, valine, leucine, aspartate, and tyrosine. Incomplete N_2_O consumption (<20% of initial dose) was observed in cultures supplemented with the aforementioned amino acid combination plus methionine. N_2_O reduction and growth of the *Desulfosporosinus* sp. occurred without delay in cultures supplied with a 15-amino acid mixture (Fig. 2 C). Omission of single amino acids from the 15-amino acid mixture led to incomplete N_2_O reduction, similar to what was observed with the 6-amino acid combination. Efforts to isolate the *Desulfosporosinus* sp. in medium without pyruvate but amended with amino acids were unsuccessful because of concomitant growth of the *Serratia* sp., as verified with qPCR.

### pH range of acidophilic N_2_O reduction by the *Desulfosporosinus* sp

Growth assays with co-culture EV were performed to determine the pH range for N_2_O reduction. Co-culture EV reduced N_2_O at pH 4.5, 5.0 and 6.0, but not at pH 3.5, 7.0 and 8.0. Cultures at pH 4.5 exhibited about two times longer (i.e., 10 versus 5 days) lag periods prior to the onset of N_2_O consumption than cultures incubated at pH 5.0 or 6.0 (*SI Appendix*, Fig. S7 A). In medium without amino acid supplementation, pyruvate fermentation is required for the initiation of N_2_O consumption, raising the question if pH impacts pyruvate fermentation by the *Serratia* sp., N_2_O reduction by the *Desulfosporosinus* sp., or both processes. Axenic *Serratia* sp. cultures fermented pyruvate over a pH range of 4.5 to 8.0, with the highest pyruvate consumption rates observed at pH 6.0 and 7.0 (1.47 ± 0.04 mmol L^-1^ day^-1^), and the lowest rates measured at pH 4.5 (0.43 ± 0.05 mmol L^-1^ day^-1^) (*SI Appendix*, Fig. S7 B). The N_2_O consumption rates in co-culture EV between pH 4.5 to 6.0 ranged from 0.24 ± 0.01 to 0.26 ± 0.01 mmol L^-1^ day^-1^ (*SI Appendix*, Fig. S7 C). Since N_2_O reduction in the absence of amino acids only commenced following pyruvate consumption (Fig. 1 C), pyruvate fermentation by *Serratia* sp., not N_2_O reduction by *Desulfosporosinus* sp., explains the extended lag periods observed at pH 4.5 (*SI Appendix*, Fig. S7 A). Consistently, shorter lag phase for both N_2_O reduction and *Desulfosporosinus* growth were observed in co-culture EV amended with the amino acid mixture (Fig. 2 C).

### Phylogenomic analysis

Phylogenomic reconstruction based on concatenated alignment of 120 bacterial marker genes corroborated the affiliation of the N_2_O-reducing bacterium with the genus *Desulfosporosinus* (Fig. 3). The genus *Desulfosporosinus* comprises strictly anaerobic, sulfate-reducing bacteria, and *Desulfosporosinus acididurans* strain SJ4 and *Desulfosporosinus acidiphilus* strain M1 were characterized as acidophilic sulfate reducers. The N_2_O-reducing *Desulfosporosinsus* sp. in co-culture EV possesses the *aprAB* and *dsrAB* genes encoding adenylyl sulfate reductase and dissimilatory sulfate reductase, respectively, but lacks the *sat* gene encoding sulfate adenylyl transferase sulfurylase (*SI Appendix*, Fig. S8 and Text S4).

**Fig. 3.**
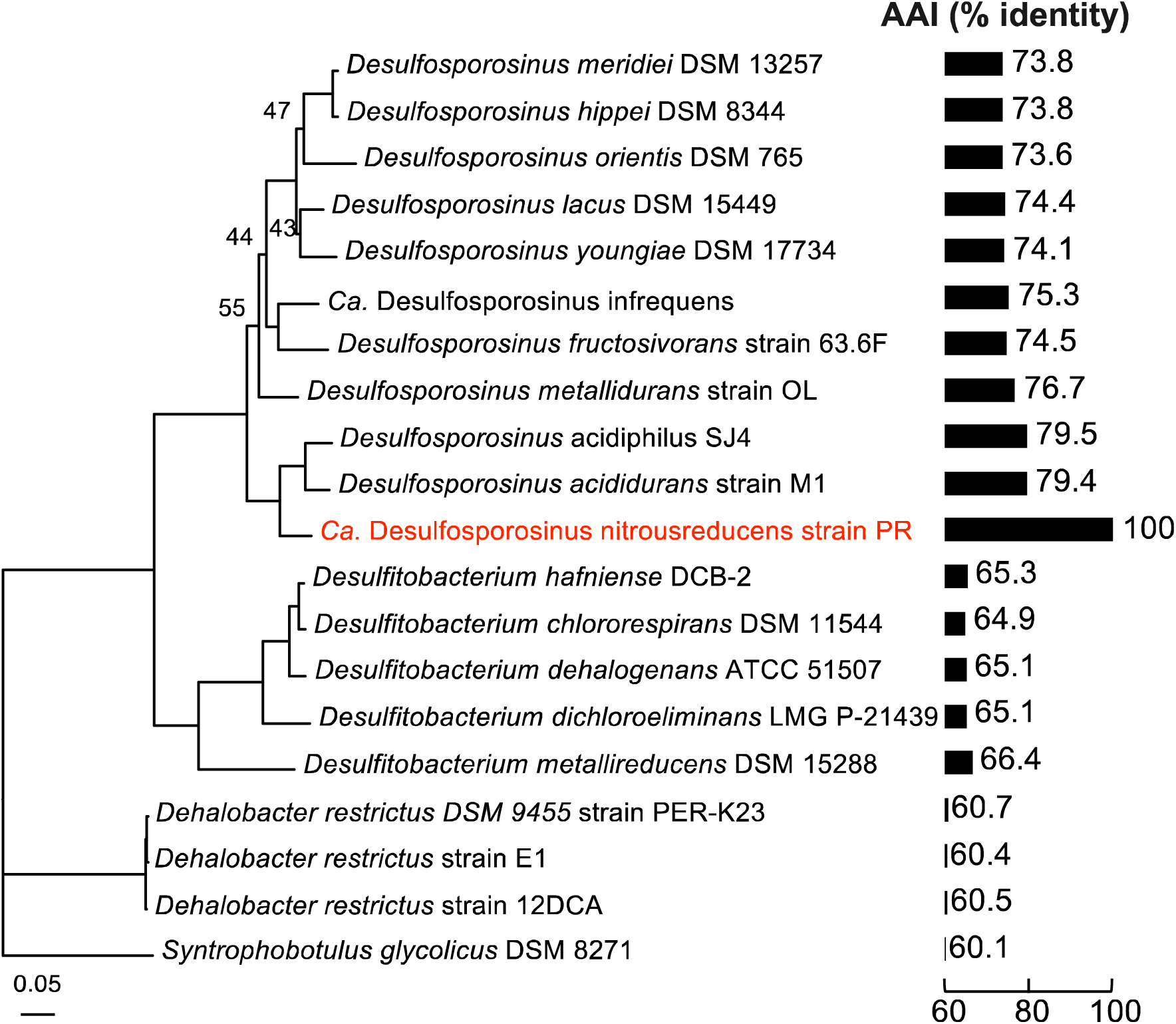
Phylogenomic and Average Amino acid Identity (AAI) analyses indicate that the N_2_O-reducing strain PR in co-culture EV represents a new species of the genus *Desulfosporosinus*. Phylogenomic analysis was based on 120 conserved marker genes and included *Peptococcaceae* genomes available from NCBI. Bootstrap values higher than 90 are not displayed. The scale bar indicates 0.05 nucleotide acid substitution per site. Bar plots display the genome-wide AAI (%) between the N_2_O-reducing ‘*Ca*. Desulfosporosinus nitrousreducens’ and related isolates with sequenced genomes.

To provide experimental evidence that the N_2_O-reducing *Desulfosporosinus* sp. in co-culture EV lacks the ability to reduce sulfate, a hallmark feature of the genus *Desulfosporosinus*, comparative growth studies were performed. The N_2_O-reducing *Desulfosporosinsus* sp. in co-culture EV did not grow with sulfate as sole electron acceptor, consistent with an incomplete dissimilatory sulfate reduction pathway (*SI Appendix*, Fig. S9 A). *Desulfosporosinus acididurans* strain D (25), a close relative of the N_2_O-reducing *Desulfosporosinus* sp. in co-culture EV, grew with sulfate in pH 5.5 medium, but did not grow with N_2_O under the same incubation conditions (*SI Appendix*, Fig. S9 B). These observations corroborate that the N_2_O-reducing *Desulfosporosinus* sp. lacks the ability to perform dissimilatory sulfate reduction. Based on phylogenetic and physiologic features, the N_2_O-reducer in culture EV represents a novel *Desulfosporosinus* sp., for which the name ‘*Candidatus* Desulfosporosinus nitrousreducens’ strain PR is proposed.

### Genetic underpinning of N_2_O reduction in ‘*Ca.* Desulfosporosinus nitrousreducens’ strain PR

The strain PR genome harbors a single *nosZ* gene affiliated with clade II *nosZ* sequences (Fig. 4). Independent branch placement of the strain PR NosZ on the clade II NosZ tree suggests an ancient divergence; a finding supported by amino acid identity (AI) value comparisons between the NosZ sequences versus their homologs in the genomes of ‘*Ca.* Desulfosporosinus nitrousreducens’ strain PR and *Desulfosporosinus meridiei* (AI: 44% versus 73.83%) (Fig. 3 and Fig. 4 B). The NosZ of ‘*Ca.* Desulfosporosinus nitrousreducens’ strain PR is more similar (AI: 45%) to the taxonomically distant relative *Desulfotomaculum ruminis*.

**Fig. 4.**
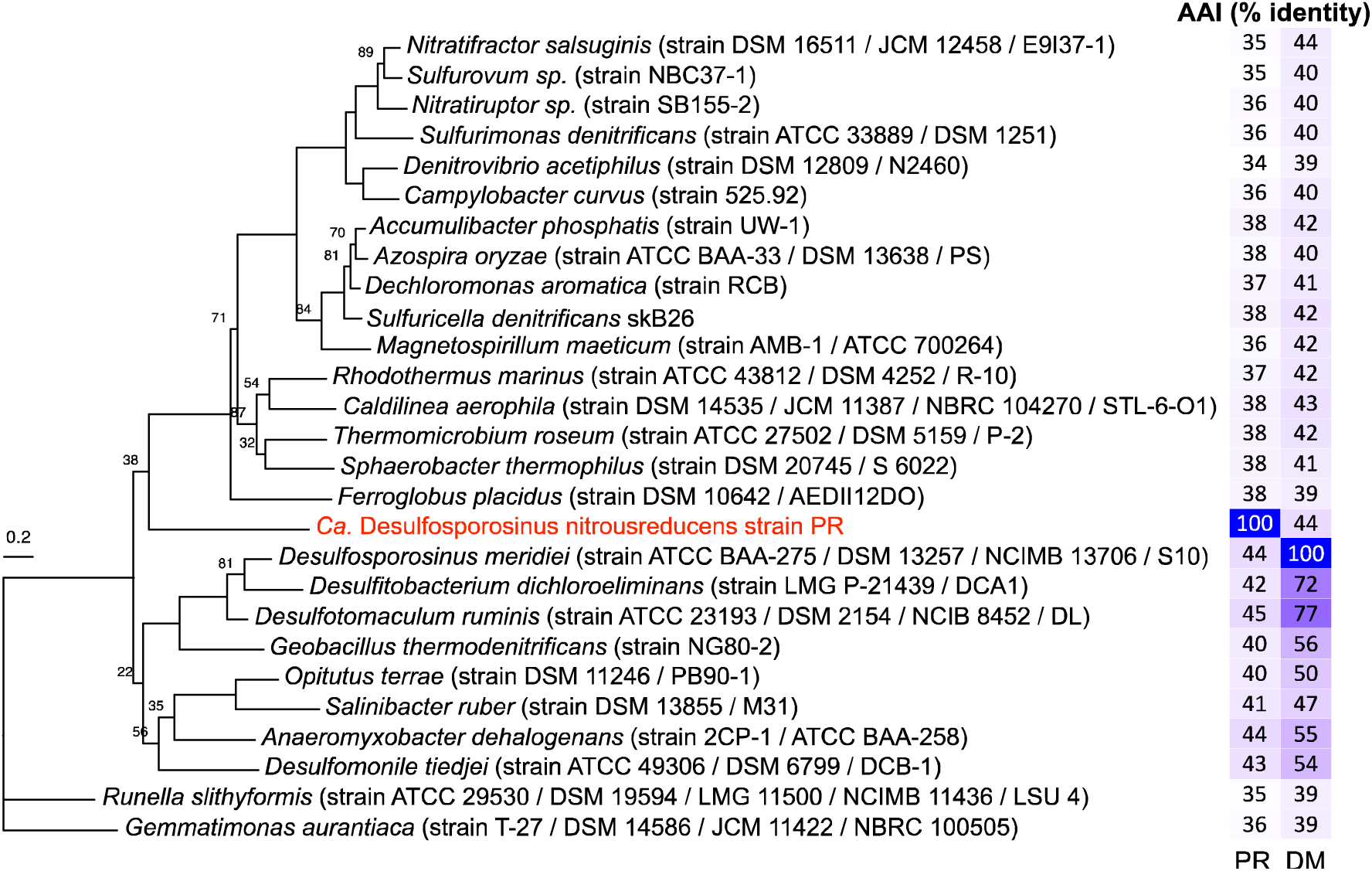
Relatedness and similarity of the clade II NosZ of ‘*Ca.* Desulfosporosinus nitrousreducens’ strain PR to representative clade II NosZ. The tree represents a phylogenetic reconstruction of select clade II NosZ protein sequences. The clade II NosZ of *Gemmatimonas aurantiaca* was used to root the tree. The scale bar indicates 0.2 amino acid substitution per site. Numbers at nodes are bootstrap values smaller than 90. The two-column heatmap shows the AAI values between the NosZ of strain PR (PR) and *Desulfosporosinus meridiei* (DM) to other clade II NosZ sequences, with the darker shades of blue indicating higher percent AAI values.

Comparative analysis of the strain PR *nos* operon with bacterial and archaeal counterparts corroborated characteristic clade II features, including a Sec translocation system, genes encoding cytochromes and an iron-sulfur protein, and a *nosB* gene located immediately downstream of *nosZ* (Fig. 5). *nosB* encodes a transmembrane protein of unknown function and has been exclusively found on clade II *nos* operons. The *nos* gene clusters found in closely related taxa (e.g., *Desulfosporosinus meridiei*, *Desulfitobacterium dichloroeliminans*, *Desulfitobacterium hafniense*) show similar organization; however, differences were observed in the *nos* gene cluster of ‘Ca. Desulfosporosinus nitrousreducens’ strain PR. Specifically, the genes encoding an iron-sulfur cluster protein and cytochromes precede *nosZ* in *Desulfosporosinus meridiei*, but are located downstream of two genes encoding proteins of unknown functions in strain PR (Fig. 5). Of note, among the microbes with *nos* operons and included in the analyses, only ‘*Ca.* Desulfosporosinus nitrousreducens’ and *Nitratiruptor labii* (26), both with a clade II *nos* operon, were experimentally validated to grow with N_2_O below pH 6 (Table S3).

**Fig. 5.**
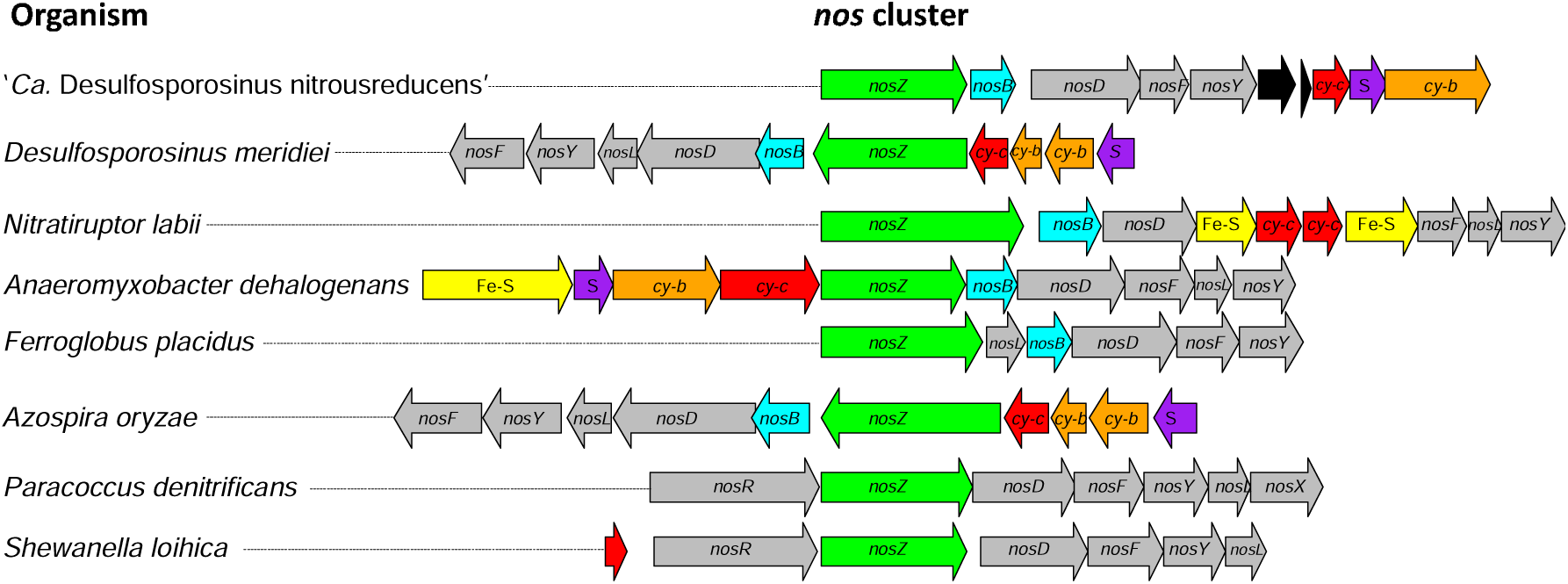
Comparison of representative *nos* clusters. Included are clade II *nos* clusters encoding select NosZ shown in Fig. 4 and select clade I *nos* clusters of bacteria with confirmed N_2_O reduction activity at circumneutral pH. The colored arrows represent genes with different functions and indicate orientation and approximate length. Green, *nosZ*; gray, *nos* accessory genes (i.e., *nosD*, *nosF*, *nosY*, *nosL*, *nosX* and *nosR*); yellow, genes encoding iron-sulfur (Fe-S) proteins; purple, genes encoding Rieske iron-sulfur proteins (S); orange (cy-b) and red (cy-c), genes encoding b-type and c-type cytochromes, respectively; cyan, *nosB* genes encoding transmembrane proteins characteristic for clade II *nos* operons; black, genes of unknown function. *Desulfosporosinus* and *Desulfitobacterium* spp., *Nitratiruptor* and *Nitratifractor* spp., and *Paracoccus* and *Bradyrhizobium* spp. share similar *nos* cluster architectures, respectively.

### Genomic insights for a commensalistic relationship

Functional annotation of the *Serratia* sp. and the ‘*Ca*. Desulfosporosinus nitrousreducens’ strain PR genomes was conducted to investigate the interspecies interactions (Fig. 6). A *ptsT* gene encoding a specific, high-affinity pyruvate/proton symporter (27) and genes implicated in pyruvate fermentation (i.e., *pflAB*, *poxB*) are present on the *Serratia* genome, but are missing on the strain PR genome, consistent with the physiological characterization results. *fdhC* genes encoding a formate transporter are present on both genomes, but only the strain PR genome harbors the *fdh* gene cluster encoding a formate dehydrogenase complex (*SI Appendix*, Fig. S8), consistent with the observation that the *Serratia* sp. excretes formate, which strain PR utilizes as electron donor for N_2_O reduction (*SI Appendix*, Fig. S1). Gene clusters encoding two different Ni/Fe-type hydrogenases (i.e., *hyp* and hya gene *clusters*) (*SI Appendix*, Fig. S8) and a complete *nos* gene cluster (Fig. 5) are present on the strain PR genome, but not on the *Serratia* sp. genome.

**Fig. 6.**
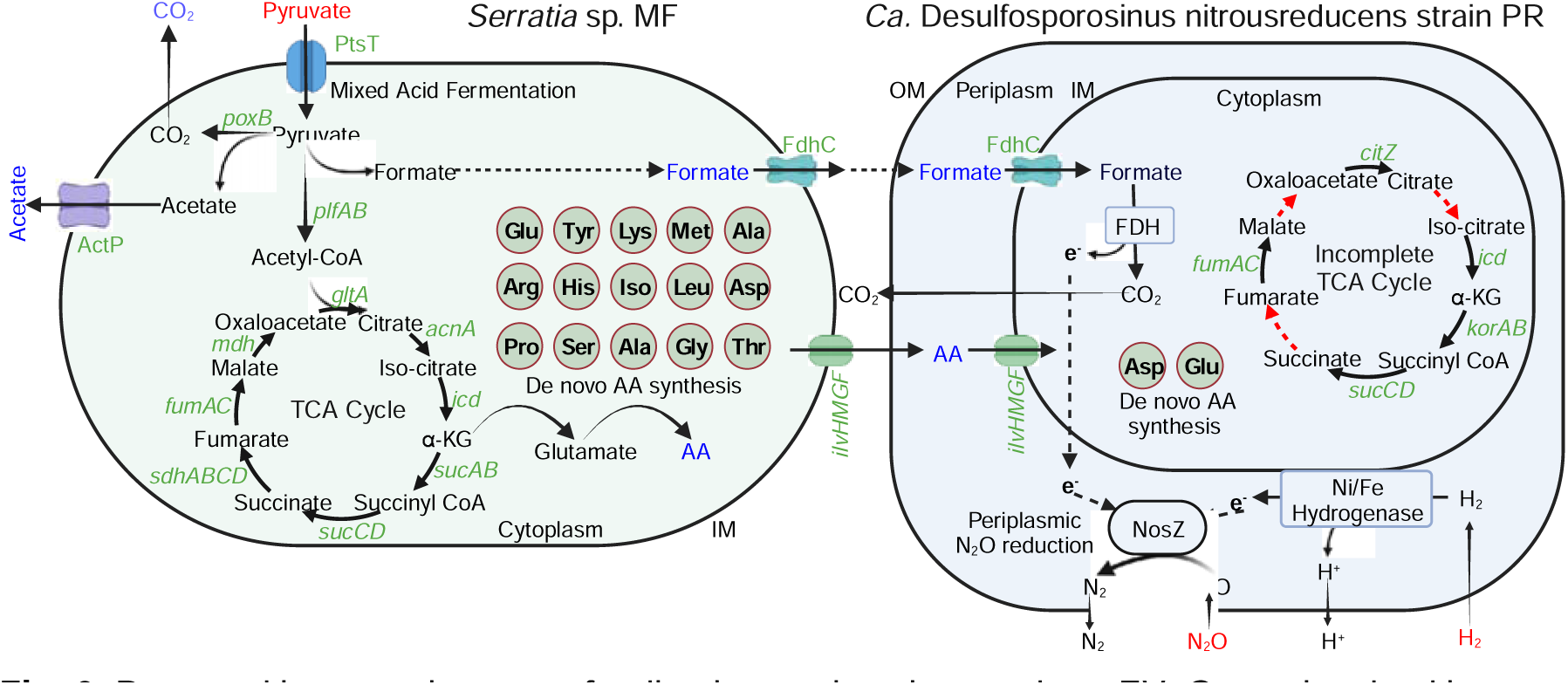
Proposed interspecies cross-feeding interactions in co-culture EV. Genes involved in mixed acid fermentation are only found on the *Serratia* sp. strain MF genome, and genes involved in periplasmic N_2_O reduction are exclusive to ‘*Ca.* Desulfosporosinus nitrousreducens’ strain PR. External substrates (i.e., pyruvate or the 15-amino acid mixture, H_2_, N_2_O) provided to co-culture EV are shown in red font, and metabolites produced by *Serratia* are shown in blue font. Fifteen versus two complete amino acid biosynthesis pathways are present on the *Serratia* sp. strain MF and ‘*Ca.* Desulfosporosinus nitrousreducens’ strain PR genomes, respectively. Strain PR has an incomplete TCA cycle, and the red dashed arrows indicate the absence of the corresponding genes. TCA cycle: tricarboxylic acid cycle; AA: amino acids; FDH: formate dehydrogenase complex; NosZ: nitrous oxide reductase; OM: outer membrane; IM: inner membrane.

Based on the KEGG (28) and Uni-Prot databases (29), the *Serratia* genome contains the biosynthesis pathways (100% completeness) for aspartate, lysine, threonine, tryptophan, isoleucine, serine, leucine, valine, glutamate, arginine, proline, methionine, tyrosine, cysteine, and histidine. In contrast, only aspartate and glutamate biosynthesis are predicted to be complete on the strain PR genome, whereas the completeness level for biosynthesis pathways of other amino acids was below 80%. The *Serratia* genome encodes a complete set of TCA cycle enzymes, indicating the potential for forming aspartate and glutamate via transamination of oxaloacetate and α-ketoglutarate. In contrast, the strain PR genome lacks genes encoding malate dehydrogenase, citrate synthase, and aconitate hydratase, indicative of an incomplete TCA cycle. Therefore, strain PR is deficient of *de novo* formation of precursors for glutamate, aspartate, alanine, and related amino acids (30). A high-affinity amino acid transport system was found on the strain PR genome (*SI Appendix*, Fig. S8), suggesting this bacterium can efficiently acquire extracellular amino acids to meet its nutritional requirements.

### Distribution of the novel N_2_O reducer in soil metagenomes

For estimating the distribution of the *Serratia* sp. and ‘*Ca*. Desulfosporosinus nitrousreducens’ strain PR in acidic and some circumneutral pH soils, fragments of soil metagenomes (n=195) from a prior study (31) were aligned to the *Serratia* sp. strain MF and the ‘*Ca*. Desulfosporosinus nitrousreducens’ strain PR genomes (*SI Appendix*, Fig. S10 A). The percentage of metagenome fragments that aligned to strain MF and strain PR ranged between 0.11-1.15% and 0.039-0.094%, respectively. RandomForest analysis revealed weak relationships between the presence of ‘*Ca*. Desulfosporosinus nitrousreducens’ strain PR and soil C/N ratio, pH, and NH ^+^ content. In contrast, *Serratia* sp. strain MF showed the strongest relationship to ‘*Ca*. Desulfosporosinus nitrousreducens’ strain PR, explaining 32.6% of the variability of strain PR abundance in the metagenomes included in the analysis (*SI Appendix*, Fig. S10 B). Furthermore, the relative abundances of the two populations showed a positive correlation (*SI Appendix*, Fig. S10 C), suggesting the observed commensalistic relationship may not have coincidentally developed in co-culture EV, and possibly occurs in soil habitats. Soil organic matter (r = -0.35, p < 0.01) and NO ^-^ (r = -0.31, p < 0.01) content strongly correlated with the presence of ‘*Ca*. Desulfosporosinus nitrousreducens’ strain PR.

## Discussion

Biological processes fix about 180 Tg N per year (32) and conventional agriculture introduces more than 100 Tg N of chemically fixed N each year (33). Biogeochemical cycling converts a substantial fraction of fixed N to N_2_O, and rising atmospheric N_2_O concentrations indicate that N_2_O sources outpace microbial N_2_O sinks. Soil acidification is a major concern relevant to N cycling and N_2_O emissions. Microbial processes (e.g., nitrification and denitrification) involved in N_2_O production have been reported to cause greater N_2_O emissions at pH < 6 than at circumneutral pH (34, 35). In contrast, characterized N_2_O reducers from soil habitats consume N_2_O within a narrow pH window ranging from 6 to 8 with generally weak activity approaching pH 6.0 and no activity at lower pH (18, 36–38). Interestingly, metagenomic studies revealed that acidic Luquillo Experimental Forest (Puerto Rico) tropical soils and temperate circumneutral pH soils harbor a comparable diversity of *nosZ* genes (22), and that pH selects for different N_2_O reducers in Luquillo tropical soil microcosms (23).

A few studies reported limited N_2_O reduction activity in acidic microcosms, but enrichment cultures for detailed experimentation were not obtained (12, 35, 39). Possible explanations for the observed N_2_O consumption in acidic microcosms include residual activity of existing N_2_O-reducing biomass, or the presence of microsites on soil particles where solid phase properties influence local pH, generating pH conditions not captured by bulk aqueous phase pH measurements (40, 41). Soil microcosms providing microsites with favorable (i.e., higher) pH conditions can give the false impression of low pH N_2_O consumption; however, removal of solids during transfers eliminates this niche and microorganisms experience bulk phase pH conditions, resulting in the cessation of N_2_O reduction activity. Our work with acidic tropical soils alludes to another crucial issue, explicitly the choice of carbon source for the successful transition from microcosms to soil-free enrichment cultures. Lactate sustained N_2_O reduction in pH 4.5 Luquillo tropical soil microcosms, but transfer cultures commenced N_2_O reduction only when pyruvate substituted lactate. Lactate has a higher p*K*_a_ value than pyruvate (3.8 versus 2.45), indicating that a larger fraction of protonated, and potentially toxic, lactic acid exists at pH 4.5 (42). As discussed above, in soil microcosms, particles with ion exchange capacity (i.e., microsites) can suppress inhibitory effects of protonated organic acids, a possible explanation why lactate supported N_2_O reduction in the microcosms but not in the enrichment cultures.

Fifteen repeated transfers with N_2_O, pyruvate, and H_2_ yielded a co-culture comprising a *Serratia* sp. and a *Desulfosporosinus* sp. The rapid enrichment of a co-culture may be surprising considering that pyruvate and H_2_ are excellent substrates for many soil microbes. N_2_O was the sole electron acceptor provided to the defined basal salt medium, with some CO_2_ being formed during pyruvate fermentation (Phase I); however, no evidence was obtained for H_2_-driven CO_2_ reduction to acetate or to methane. N_2_O has been recognized as potent inhibitor of corrinoid-dependent enzyme systems (9, 10), and both CO_2_/H_2_ reductive acetogenesis and hydrogenotrophic methanogenesis would not be expected to occur in the enrichment cultures, a prediction supported by the analytical measurements.

Available axenic and mixed cultures obtained from circumneutral pH soils reduce N_2_O at circumneutral pH, but not below pH 6.0 (15, 36, 43). The discovery and cultivation of co-culture EV comprising ‘*Ca.* Desulfosporosinus nitrousreducens’ strain PR extends the pH range for microbial growth with N_2_O as electron acceptor to pH 4.5. Metagenomic studies routinely detect *nosZ* in acidic soils providing evidence that bacteria capable of low pH N_2_O reduction exists and are likely more widely distributed in acidic soil habitats (22, 44, 45). The distribution of bacteria capable of low pH N_2_O reduction may extend to the deep marine biosphere, a hypothesis supported by the recent discovery of the thermophile *Nitratiruptor labii* strain HRV44 with the capacity to reduce N_2_O over a pH range of 5.4 to 6.4 (26).

‘*Ca.* Desulfosporosinus nitrousreducens’ strain PR reduces N_2_O between pH 4.5 and 6.0, with no N_2_O reduction observed at or above pH 7. This finding implies that ‘*Ca.* Desulfosporosinus nitrousreducens’ cannot be enriched with N_2_O as electron acceptor at or above pH 6.5, suggesting the maintenance of acidic pH conditions during enrichment is crucial to cultivate new microorganisms capable of low pH N_2_O reduction. Available N_2_O-reducing soil isolates enriched and isolated at circumneutral pH cannot grow with N_2_O as electron acceptor below pH 6.0.

Apparently, pH selects for distinct groups of N_2_O reducers, with prior research mostly focused on soil isolates obtained at circumneutral pH. The discovery of ‘*Ca.* Desulfosporosinus nitrousreducens’ strain PR lends credibility to the hypothesis that the diverse *nosZ* genes found in metagenomes of acidic soils (22) may indeed be functional. Further, the cultivation of strain PR provides a blueprint for unraveling the unknown microbiology of low pH N_2_O reduction and exploring the geochemical parameters that govern this process in acidic soils.

‘*Ca.* Desulfosporosinus nitrousreducens’ strain PR possesses a clade II *nos* operon similar to those found in clade II neutrophilic N_2_O reducers without distinguishing features based on gene content and *nos* operon structure (Fig. 5). Experimental work with *Paracoccus denitrificans*, a model organism harboring a clade I *nosZ* and used for studying denitrification, has shown that acidic pH impairs N_2_O reductase assembly and maturation (18). NosZ is a periplasmic enzyme with a distinguishing feature of clade I versus clade II NosZ being the mechanism of protein transport across the cytoplasmic membrane. Clade II NosZ follows the general Secretion route known as Sec-pathway, which translocates unfolded proteins. In contrast, clade I NosZ are translocated in a folded state via the Twin-arginine pathway (Tat-pathway) (46). Differences exist between clade I and cade II *nos* operons, and *nosB*, encoding a transmembrane protein of unknown function, has been exclusively found associated with clade II *nos* clusters (Fig. 5) (47, 48). To what extent gene content and the secretion pathway influence the pH range of N_2_O reduction is unclear and warrants further genetic/biochemical studies. Other factors that can impact N_2_O reduction at acidic pH include an organism’s ability to cope with the potential toxicity of protonated organic acids and maintain pH homeostasis (49, 50). The ‘*Ca.* Desulfosporosinus nitrousreducens’ strain PR genome harbors multiple genes associated with DNA repair and potassium transport, suggesting this bacterium can respond to pH stress.

Soils harbor diverse microbiomes with intricate interaction networks that govern soil biogeochemical processes, including N_2_O turnover (51), and define the functional dynamics of microbiomes (52–54). Interspecies cooperation between bacteria can enhance N_2_O reduction via promoting electron transfer (55), the provision of essential nutrients (as demonstrated in co-culture EV), or limit N_2_O reduction due to competition for electron donor(s) or metal cofactors (i.e., copper) (56, 57). Metabolomic workflows unraveled that *Serratia* sp. strain MF excretes amino acids during growth with pyruvate, which ‘*Ca.* Desulfosporosinus nitrousreducens’ strain PR requires to initiate N_2_O reduction, a finding supported by genome functional predictions (i.e., 15 complete AA biosynthesis pathways in strain MF versus only two complete AA biosynthesis pathways in strain PR). An amino acid mixture supported N_2_O reduction by strain PR in the absence of pyruvate; however, slight growth of *Serratia* sp. strain MF was observed, preventing the isolation of the N_2_O reducer Interspecies interactions based on amino acid auxotrophies have been implicated in shaping dynamic anaerobic microbial communities, bolster community resilience, and thus promote functional stability (52). Other microbes can potentially fulfill the nutritional demands of ‘*Ca.* Desulfosporosinus nitrousreducens’, and the observed interrelationship between the *Serratia* sp. and strain PR might have developed coincidentally during the enrichment process; however, the co-occurrence pattern does imply some level of specificity and that the interactions may occur in situ.

Members of the genus *Desulfosporosinus* have been characterized as strictly anaerobic sulfate reducers with the capacity to grow autotrophically with H_2_, CO_2_, and sulfate, or, in the absence of sulfate, with pyruvate (58). Most characterized *Desulfosporosinus* spp. show optimum growth at circumneutral pH (∼7) conditions, except for the acidophilic isolates *Desulfosporosinus metallidurans, Desulfosporosinus acidiphilus*, *Desulfosporosinus acididurans*, and *Desulfosporosinus* sp. strain I2, which perform sulfate reduction at pH 4.0, 3.6, 3.8, and 2.6, respectively (25, 42, 59–61). Among the 10 *Desulfosporosinus* species with sequenced genomes, only *Desulfosporosinus meridiei* DSM 13257 carries a *nos* cluster (47). *Desulfosporosinus meridiei* DSM 13257 grows within a narrow range of pH 6-8 (58), and its ability to reduce N_2_O has not been demonstrated. ‘*Ca.* Desulfosporosinus nitrousreducens’ strain PR lacks the hallmark feature of sulfate reduction and is the first acidophilic soil bacterium capable of growth with N_2_O as electron acceptor at pH 4.5, but not at pH 7 or above. Strain PR couples N_2_O reduction and growth at pH 4.5 with the oxidation of H_2_ or formate, and our experimental efforts with co-culture EV could not demonstrate the utilization of other electron donors. The four characterized acidophilic representatives of the genus *Desulfosporosinus* show considerable versatility, and various organic acids, alcohols, and sugars, in addition to H_2_, support sulfate reduction (25, 59, 60). The utilization of H_2_ as electron donor appears to be a shared feature of *Desulfosporosinus* spp., and two or more gene clusters encoding hydrogenase complexes were found on available *Desulfosporosinus* spp. genomes (61–63).

Escalating usage of N fertilizers to meet societal demands for agricultural products increases N_2_O emissions due to accelerated N cycling and associated N_2_O formation and due to soil acidification, which presumably slows N_2_O consumption. Liming is commonly employed to ameliorate soil acidity, a practice considered beneficial for curbing N_2_O emissions based on the assumption that microbial N_2_O reduction is limited to circumneutral pH soils (20, 35, 36, 64). Our findings demonstrate that soils harbors microorganisms (e.g., ‘*Ca.* Desulfosporosinus nitrousreducens’ strain PR) that utilize N_2_O as growth-supporting electron acceptor between pH 4.5 and 6.5.

Apparently, N_2_O consumption is not limited to circumneutral pH conditions and acidophilic respiratory N_2_O reducers exist in acidic soils. Thus, increased N_2_O production due to fertilizer application could potentially be mitigated, at least partially, by microbial N_2_O consumption in acidic soils. Recent efforts have shown substantial N_2_O emission reductions from soils treated with biofertilizers or augmented with mixed cultures harboring clade II N_2_O reducers (65, 66). Based on metagenomic surveys, bacteria capable of low pH N_2_O are not limited to acidic tropical soils, and are more broadly distributed in terrestrial ecosystems (23). The discovery of a naturally occurring soil bacterium that couples N_2_O consumption to growth under acidic pH conditions offers opportunities for the knowledge-based manipulation of agricultural soils to stimulate microbial N_2_O consumption. Curbing undesirable N_2_O emissions at the field scale would allow farmers to further reduce their greenhouse gas emissions footprint and potentially earn carbon credits.

## Materials and Methods

Details of all methods used in this study are described in SI Appendix and SI Materials and Methods. SI Materials and Methods include details on (i) sample collection; (ii) experimental approaches including establishment of the enrichment culture, isolation efforts, and physiological characterization; (iii) molecular techniques including DNA extraction, qPCR, sequencing, and metabolomic workflows; and (iv) computational analyses and methodologies. Additional SI References provide information about procedures and analytical techniques.

## Data availability

All sequencing data generated and used in this study are available in the NCBI database, with all accession numbers summarized in Table S2.

## Code availability

Code used for data processing and the production of figures is publicly available on GitHub at https://github.com/utguang/Paper-code.

## Author Contributions

G.H. and F.E.L. conceptualized research and designed experiments. G.H performed experimental work, including cultivation, phenotypic and molecular characterization, and bioinformatic analysis. G.C, Y.X, and C.S. provided technical expertise. K.T.K. provided guidance on bioinformatic analyses. All authors contributed to data analysis and interpretation, and G.H. and F.E.L. wrote the manuscript with inputs from Y.X., G.C., K.T.K., and M.A.R.

## Competing Interest Statement

The authors have no competing interests.

## Classification

BIOLOGICAL SCIENCES, Microbiology.

## Acknowledgments

The authors acknowledge funding through the Dimensions of Biodiversity program of the US National Science Foundation (awards 1831599 to F.E.L. and 1831582 to K.T.K.). GH acknowledges support from the China Scholarship Council. DNA sequencing and metabolome analysis were performed by the University of Tennessee Genomics Core and the Biological and Small Molecule Mass Spectrometry Core, respectively. We thank Dr. Irene Sánchez-Andrea, Wageningen University & Research, for making available a culture of *Desulfosporosinus acididurans* strain D for comparative growth studies, and Dr. Yanchen Sun for providing the N_2_O-reducing tropical soil microcosm.

## Notes

### Competing Interest Statement

The authors have declared no competing interest.

