## Supplementary material for "Sustained bacterial N_2_O reduction at acidic pH": SI Appendix

**Corresponding author:** Frank E. Löffler

**This PDF file includes:**

Supporting text

Figures S1 to S10

Tables S1, S2, S3, S4, S6 and S7

SI References

Supporting Information Text

**Materials and Methods**

**Soil sampling locations and microcosms.** Soil samples were collected in August 2018 at the El Verde research station in the El Yunque Natural Forest in Puerto Rico (1). The measured soil pH was 4.45 and characteristic for the region. Vertical distance of the El Verde research station to mean sea level is 434 meters. Fresh soil materials from 9 to 18 cm depth were used to establish pH 4.5 laboratory microcosm that were amended with N_2_O and lactate (2).

**Enrichment process.** Transfer cultures were established in 160-mL glass serum bottles containing 100 mL of anoxic, completely synthetic, defined basal salt medium prepared as described (3) with the following modifications. NaHCO_3_ and Na_2_S were omitted and the total KH_2_PO_4_ concentration was increased to 7.0 g L^-1^. l-cysteine (0.2 mM) was the sole reductant unless indicated otherwise. The medium pH was 4.27 to 4.35. The bottles with N_2_ headspace were sealed with butyl rubber stoppers (Bellco Glass, Vineland, NJ, USA) held in place with aluminum crimp caps and autoclaved. All subsequent amendments to the medium bottles used sterile plastic syringes and needles to augment the medium with aqueous filter-sterilized (0.2 µm polyethersulfone membrane filter, Thermo Scientific, Waltham, MA, USA) stock solutions and undiluted gases (4). Ten mL of N_2_O gas (416 µmol, 4.16 mM nominal; 99.5%) was added 24 hours prior to inoculation from an El Verde microcosm showing N_2_O reduction activity. The microcosm was manually shaken before 1 mL aliquots were transferred with a 3-mL plastic syringe and a 2-gauge needle. Initial attempts to obtain solids-free enrichment cultures with 5 mM lactate as carbon source and electron donor showed no N_2_O reduction. As such, the following substrates were subsequently tested in the transfer cultures: 5 mM propionate, 20 mM pyruvate, 20 mM pyruvate plus 10 mL (416 µmol, 4.16 mM nominal) hydrogen (H_2_), 1 mM formate plus 1 mM acetate and 5 mL (208 µmol, 2.08 mM nominal) CO_2_, and 0.1 or 10 g L^-1^ yeast extract. Subsequent transfers (3%, v/v) used medium supplemented with 0.5 or 2.5 mM pyruvate and 10 mL H_2_, and occurred when the initial dose of 10 mL N_2_O had been consumed. All culture vessels were incubated in upright position at 30ºC in the dark without agitation.

**Microbial community analysis.** 16S rRNA gene amplicon sequencing was performed with samples collected from 6^th^-generation transfer culture following complete N_2_O consumption, and 9^th^-generation transfer cultures following complete pyruvate consumption (Phase I) and complete N_2_O consumption (Phase II). Biomass from 1 mL of culture suspension samples were collected by centrifugation (10,000 x g, 20 min, 4 ºC), and genomic DNA was isolated from the cell pellets using the DNeasy PowerSoil Kit (Qiagen, Hilden, Germany). 16S rRNA gene-based amplicon sequencing was conducted at the University of Tennessee Genomics Core following published procedures (5). Primer sets used in amplicon sequencing of 6^th^ and 9^th^ transfer cultures are 341F-785R and 515F-805R, respectively (6).

Analysis of amplicon reads was conducted with nf-core/ampliseq v2.3.1 using Nextflow (7). Software used in nf-core/ampliseq was containerized with Singularity v3.8.6 (8). Amplicon read quality was evaluated with FastQC v0.11.9 (9) and primer removal used Cutadapt v3.4 (10). Quality control including removal of sequences with poor quality, denoising, and chimera removal was performed, and amplicon sequence variants (ASVs) were inferred using DADA2 (11). Barrnap v0.9 was used to discriminate rRNA sequences as potential contamination (12). ASVs were taxonomically classified based on the Silva v138.1 database (13). Relative and absolute abundances of ASVs were calculated using Qiime2 v2021.8.0 (14). Short read fragments of the El Verde soil metagenome representing 16S rRNA genes were identified and extracted using Parallel-Meta Suite v3.7 (15).

**Isolation and classification.** Following 15 consecutive transfers, 100 µL cell suspension aliquots were serially diluted and plated on tryptic soy agar (TSA, MilliporeSigma, Maryland, US) medium. Colonies with uniform morphology were observed, and a single colony was transferred to a new TSA plate. This process was repeated three times before a single colony was transferred to liquid basal salt medium (pH 4.5) amended with 2.5 mM pyruvate, 416 µmol N_2_O, and 416 µmol H_2_. Following growth, DNA was extracted for PCR amplification with general bacterial 16S rRNA gene-targeted primer pair 8F-1541R (16), and Sanger sequencing of both strands yielded a 1,471-bp long 16S rRNA gene fragment.

Efforts to isolate the N_2_O reducer applied the dilution-to-extinction principle (3). Ten-fold dilution-to-extinction series used 20 mL glass vials containing 9 mL of basal salt medium and 0.6% (w/v) low melting agarose (MP Biomedicals, LLC., Solon, OH) with a gelling temperature below 30º were established as described (3). Each glass vial received 2.5 mM pyruvate, 1 mL (41.6 µmol, 4.16 mM nominal) H_2_ and 1 mL (41.6 µmol, 4.16 mM nominal) N_2_O following heat sterilization. Parallel 10^-1^ to 10^-10^ dilution-to-extinction series were established in liquid basal salt medium without low melting agarose, which were used to inoculate the respective soft agar dilution vials. Additional attempts to isolate the N_2_O reducer used solidified (1.5% agar, w/v) basal salt medium. A 1-mL sample of a 15^th^-generation transfer culture that actively reduced N_2_O was 10-fold serially diluted in liquid basal salt medium, and 100 µL of cell suspension aliquots were evenly distributed on the agar surface. The plates were incubated under an atmosphere of N_2_/H_2_/N_2_O (8/1/1, v/v/v), and colony formation was monitored every 2 weeks over a 6-month period.

**Quantitative PCR (qPCR).** SYBR Green qPCR assays targeting the 16S rRNA gene of the *Serratia* sp., and TaqMan qPCR assays targeting the 16S rRNA gene of the *Desulfosporosinus* sp. were designed using Geneious Prime (Table S1). Probe and primer specificities were examined by *in silico* analysis using the Primer-BLAST tool (17), and experimentally confirmed using 1,538 bp- and 1,467 bp-long synthesized linear DNA of the respective complete 16S rRNA genes of the *Serratia* sp. and the *Desulfosporosinus* sp., respectively (Integrated DNA Technologies, USA). For enumeration of *Serratia* 16S rRNA genes, 25 µL qPCR tubes received 10 µL 1X Power SYBR Green, 9.88 µL UltraPure nuclease-free water (Invitrogen, Carlsbad, CA, USA), 300 nM of each primer, and 2 µL template DNA. For enumeration of *Desulfosporosinus* 16S rRNA genes, the qPCR tubes received 10 µL TaqMan Universal PCR Master Mix (Life Technologies, Carlsbad, CA, USA), 300 nM of TaqMan probe (5’-6FAM-AAGCTGTGAAGTGGAGCCAATC-MGB-3’), 300 nM of each primer, and 2 µL template DNA (18). All qPCR assays were performed using an Applied Biosystems ViiA 7 system (Applied Biosystems, Waltham, MA, USA) with the following amplification conditions: 2 min at 50ºC and 10 min at 95ºC, followed by 40 cycles of 15 s at 95ºC and 1 min at 60ºC. The standard curves were generated using 10-fold serial dilutions of the linear DNA fragments carrying a complete sequence of the *Serratia* sp. (1,538 bp) or the *Desulfosporosinus* sp. (1,467 bp) 16S rRNA gene, covering the 70- and 72-bp qPCR target regions, respectively.

The qPCR standard curves established with the linear DNA fragments carrying complete *Serratia* sp. or *Desulfosporosinus* sp. 16S rRNA genes had slopes of -3.82 and -3.404, y-intercepts of 37.408 and 34.181, R^2^ values of 0.999 and 1, and qPCR amplification efficiencies of 82.7% and 96.7%, respectively. The linear range spanned 1.09 to 1.09 x 10^8^ gene copies per reaction with a calculated detection limit of 10.9 gene copies per reaction. The genome analysis indicated that both the *Serratia* sp. and the *Desulfosporosinus* sp. genomes carry a single 16S rRNA gene, indicating that the enumeration of 16S rRNA gene allows estimates of cell abundances. The 16S rRNA gene sequences of the *Serratia* sp. and the *Desulfosporosinus* sp. are available under NCBI accession numbers OR076433 and OR076434, respectively.

**Nutritional interactions in the co-culture.** To explore the nutritional requirements of the *Desulfosporosinus* sp., a time series metabolome analysis of culture supernatant was conducted. Briefly, the axenic *Serratia* sp. culture was grown in basal salt medium amended with 2.5 mM pyruvate, 4.16 mM (nominal) H_2_, and 4.16 mM (nominal) N_2_O. Following a 7-day incubation period, during which pyruvate was completely consumed, the bottles received 1% (v/v) co-culture EV inoculum. Cell suspension aliquots (1.5 mL) were collected and centrifuged, and the resulting cell-free supernatants were transferred to 2 mL plastic tubes and immediately stored at –80 ºC for metabolome analysis. Additional samples assessed the metabolome associated with supernatant of axenic *Serratia* sp. cultures that received 1 mM DTT instead of l-cysteine as reductant. The results of the metabolome analysis guided additional cultivation experiments with amino acid mixtures replacing pyruvate. The 100-fold concentrated, aqueous 15-amino acid stock solution contained (g L^-1^): alanine (0.5); aspartic acid (1); proline (1); tyrosine (0.3); histidine (0.3); tryptophan (0.2); arginine (0.5); isoleucine (0.5); methionine (0.4); glycine (0.3); threonine (0.5); valine (0.9); lysine (1); glutamate (1); serine (0.8). The aqueous stock solution was filter-sterilized and stored in the dark.

**Metagenome sequencing.** DNA was isolated from the axenic *Serratia* sp. culture grown with 2.5 mM pyruvate, and the N_2_O-reducing 15^th^ generation co-culture EV grown on H_2_, N_2_O, and the amino acid mixture. Metagenome sequencing was performed at the University of Tennessee Genomics Core using the Illumina NovaSeq 6000 platform. Shotgun sequencing generated a total of 494 and 387 Gbp of raw sequences from the axenic *Serratia* sp. culture and co-culture EV. Metagenomic short reads were processed using the nf-core/mag pipeline (19). Short read quality was evaluated with FastQC v0.11.9, followed by quality filtering and Illumina adapter removal using fastp v0.20.1 (20). Short reads mapped to the PhiX genome (GCA_002596845.1, ASM259684v1) with Bowtie2 v2.4.2 were removed (21). Assembly of processed short reads used Megahit2 v1.2.9 (22). Binning of assembled contigs was conducted with MetaBAT2 v2.15 based on the sequencing depth (23), and metagenome-assembled genomes (MAGs) that passed CheckM (24) were selected for further analysis. Protein-coding sequences on MAGs were predicted using MetaGeneMark-2 (25) and functional annotation used Blastp (26) against the Swiss-Prot database (27), KEGG (28) and the RAST server (29). Amino acid biosynthesis completeness was evaluated using KofamKOALA (28).

Metagenomic datasets of El Verde soil and a 15^th^ transfer culture were searched against the ‘*Ca*. Desulfosporosinus nitrousreducens’ strain PR MAG using blastn (26). The best hits were extracted using an in-house script embedded in Enveomics Collection tools (30). Graphical representation showing short reads recruited to the ‘*Ca.* Desulfosporosinus nitrousreducens’ strain PR MAG was generated with BlasTab.recplot2.R. The coverage evenness was assessed based on distribution of high nucleotide identity reads across the reference genome sequences. Nonpareil v3.4.1 using the weighted NL2SOL algorithm was used to estimate the average coverage level of the metagenomic datasets (31). Metagenome data of the original El Verde soil was downloaded from NCBI (accession number PRJEB26500). Metagenomic datasets of co-culture EV and axenic *Serratia* isolate were deposited at NCBI under accession numbers SRR24709127 and SRR24709126, respectively (Table S2).

**Comparative analysis of *nos* gene clusters.** Available genomes of select N_2_O reducers were downloaded from NCBI (Table S3). Functional annotation of the genomes was conducted using the RAST server. Transmembrane topology of the protein encoded by *nosB*, a gene located immediately adjacent to *nosZ* was confirmed using DeepTMHMM (32). Accessory genes associated with NosZ functionality were selected from a set of seven genomes and used as PSI-BLAST database to query accessory genes on contigs predicted to harbor *nosZ* on the ‘*Ca.* Desulfosporosinus nitrousreducens’ strain PR genome. The *nos* gene clusters were visualized using the gggenes package (<https://wilkox.org/gggenes/index.html>).

**Phylogenomic analysis.** Phylogenomic reconstruction was performed with genomes of the *Desulfitobacteriaceae* family available in the NCBI database (Table S4). Conserved marker genes of the 20 genomes were identified and aligned with GTDB-TK (33). Phylogenetic relationships were inferred based on the alignment of 120 concatenated bacterial marker genes using RAxML-NG (34) with 1,000 bootstrap replicates. A best fit evolutionary model was selected based on the result of Modeltest-NG (35). Calculation of average amino acid identity (AAI) and hierarchical clustering of taxa based on AAI values were conducted with EzAAI (36). Tree annotation and visualization were performed with the ggtree package (37).

**NosZ phylogenetic analysis.** NosZ reference sequences were downloaded from pre-compiled models of ROCker (38). The NosZ sequence of ‘Ca. Desulfosporosinus nitrousreducens’ strain PR was aligned to the NosZ reference sequences using MAFFT (39), and a maximum likelihood tree was created with RAxML-NG based on the best model from Modeltest-NG. The inferred tree and amino acid identity between ‘*Ca.* Desulfosporosinus nitrousreducens’ strain PR, *Desulfosporosinus meridiei* and the NosZ reference sequences were visualized using ggtree package.

**Metabolome analysis.** Cell-free samples were prepared as described (40). Briefly, 1.5 mL of 0.1 M formic acid in 4:4:2 (v:v:v) acetonitrile:water:methanol was added to 100 µL aliquots of supernatant samples. The tubes were shaken at 4°C for 20 minutes and centrifuged at 16,200 x g for 5 minutes. The supernatant was collected and dried under a steady stream of N_2_. The dried extracts were suspended in 300 µL of water prior to analysis. For water soluble metabolites, the mass analysis was performed in untargeted mode (41). The chromatographic separations utilized a Synergi 2.6 µm Hydro RP column (100 Å, 100 mm x 2.1 mm; Phenomenex, Torrance, CA) with tributylamine as an ion pairing reagent, an UltiMate 3000 binary pump (Thermo Scientific, San Jose, CA), and previously described elution conditions (40). The mass analysis was carried out using an Exactive Plus Orbitrap MS (Thermo Scientific) using negative electrospray ionization and full scan mode. Following the analysis, metabolites were identified using exact masses and retention times, and the areas under the curves (AUC) for each chromatographic peak were integrated using the open-source software package Metabolomic Analysis and Visualization Engine (41, 42). Dynamic changes of metabolites over time were assessed by comparative analysis of AUC values.

**Phenotypic characterization of co-culture EV.** To test for autotrophic growth of co-culture EV, pyruvate was replaced by 5 mL (2.08 mM nominal) of CO_2_ (99.5% purity). All experiments used triplicate cultures, and vessels without pyruvate, without H_2_, without N_2_O, or without inoculum served as controls. Growth experiments were conducted to determine the responses of the *Serratia* sp. and the *Desulfosporosinus* sp. to pH. Desired medium pH values of 3.5, 4.5, 5, 6, 7 and 8 were achieved by adjusting the mixing ratios of KH_2_PO_4_ and K_2_HPO_4_. To achieve pH 3.5, the pH 4.5 medium was adjusted with 5 M hydrochloric acid. The medium with different pH, 3.5, 4.5, 5, 6, 7 and 8, was achieved by adjusting the mix ratios of KH_2_PO_4_ and K_2_HPO_4_. Replicate incubation vessels received 4.16 mM (nominal) N_2_O and 4.16 mM H_2_ (nominal), and 2.5 mM pyruvate, following an overnight equilibration period, 1% inocula from the axenic *Serratia* sp. culture or the N_2_O-reducing co-culture EV, both pregrown in pH 4.5 medium. The replicate cultures inoculated with the *Serratia* sp. were incubated for 14-days, after which three vessels received an inoculum of co-culture EV (1%) to initiate N_2_O consumption. Three *Serratia* sp. cultures not receiving a co-culture EV inoculum served as controls. Consumption rates of pyruvate and N_2_O were calculated based on data points representing linear ranges of consumption according to

$V=\frac{N}{T_{1}-T_{0}}$	Equation 1

where V represent the consumption rate; N represent the initial amounts of pyruvate or N_2_O. T_1_ refers to timepoints when pyruvate or N_2_O were completely consumed. T_0_ for pyruvate consumption refers to day zero (i.e., after inoculation with axenic *Serratia* sp.). T_0_ for N_2_O consumption refers to day 14 (i.e., after inoculation with co-culture EV), following which a linear N_2_O consumption was observed.

**Analytical procedures.** N_2_O, CO_2_, and H_2_ were analyzed by manually injecting 100 µL headspace samples manually injected into an Agilent 3000A Micro-Gas Chromatograph (Palo Alto, CA, USA) equipped with Plot Q and molecular sieve columns coupled with a thermal conductivity detector as described (43). Aqueous concentrations (µM) were calculated from the headspace partial pressures based on reported dimensionless Henry’s law constants for N_2_O (2.4×10^-4^), H_2_ (7.8×10^-6^) and CO_2_ (3.3×10^-4^) (44) according to

$HRT=\frac{C_{g}}{C_{aq}}$	Equation 2

Where *H* = Henry’s law constants, *R* = universal gas constant, *T* = temperature, *C_g_* = headspace gas-phase concentration and *C_aq_* = liquid-phase (dissolved) concentration. Five-point standard curves for N_2_O, CO_2_ and H_2_ spanned concentration ranges of 8,333 to 133,333 ppmv. Pyruvate, acetate and formate were measured with an Agilent 1200 Series high-performance liquid chromatography (HPLC) system (Palo Alto, CA, USA) as described (43). pH was measured in 0.4 mL samples of culture supernatant following removal of cells by centrifugation with a pH electrode.

**Supplementary Text S1: Phenotypic characterization of the N_2_O-reducing culture.** Cultivation experiments were performed in 160 mL glass serum bottles containing 100 mL of defined basal salt medium. The consortium consumed 239 ± 1.16 µmol pyruvate within 7 days (Phase I) in the presence or absence of N_2_O, and acetate (136 ± 6.05 µmol), formate (58.5 ± 3.33 µmol) and CO_2_ (201 ± 5.53 µmol) were produced (Fig. S1 A). In cultures without N_2_O, formate was stable but was readily consumed when cultures were supplemented with N_2_O (Fig. S1 A, C, E). Measurable acetate consumption did not occur in any of the vessels indicating that acetate is not an electron donor for N_2_O reduction (Fig. S1 A, C, E).

In vessels that received pyruvate, H_2_, and N_2_O, H_2_ and N_2_O were simultaneously consumed following the depletion of pyruvate (day 10, Fig. S1 B). N_2_O consumption (Phase II) continued until H_2_ became limiting but resumed when additional H_2_ was provided (day 33, Fig. S1 B). In cultures lacking N_2_O, H_2_ was stable, but consumption commenced immediately following the addition of N_2_O (Fig. S1 D). In replicate vessels without H_2_, N_2_O was consumed after day 10, apparently coupled to the oxidation of formate, a product of pyruvate fermentation (Fig. S1 E, F). Pyruvate consumption (Phase I) and the production of formate, acetate and CO_2_ as fermentation products preceded measurable consumption of N_2_O and H_2_ (or formate) (Phase II), suggesting that pyruvate was not a direct electron donor for N_2_O reduction (Fig. S1 A-F). Following an 18-day incubation period, triplicate cultures that received pyruvate (239 ± 1.16 µmol), H_2_ (390 ± 1.66 µmol), and N_2_O (378 ± 11.7 µmol) had completely consumed N_2_O and nearly half of the exogenously added H_2_ (143 ± 8.96 µmol) (Fig. S1 B). Reduction of one molecule N_2_O requires two electrons derived from the oxidation of one molecule of H_2_ (i.e., H_2_ + N_2_O → H_2_O + N_2_). The amount of H_2_ oxidized did not reach the expected 1:1 stoichiometry, indicating that products from pyruvate fermentation (e.g., formate) serve as electron donors for N_2_O reduction. Consistently, in cultures that received a low amount of pyruvate (i.e., 50 µmol), H_2_ oxidation (361.2 ± 5.68 µmol) closely matched the amount of N_2_O reduced (377 ± 3.53 µmol) (Fig. S1 G). Pyruvate, H_2_, and N_2_O consumption were not apparent in vessels without inoculum (Fig. S1 H). No growth occurred in cultures where CO_2_ replaced pyruvate, suggesting no autotrophic activity (Fig. S1 I).

**Supplementary Text S2: Consecutive transfers with N_2_O as electron acceptor effectively enrich an N_2_O-reducing consortium.** Shotgun metagenome sequencing performed on the original El Verde soil recovered 2,718 short read fragments (150 bp) representing 16S rRNA genes, which could be assigned to 187 bacterial genera (1). 16S rRNA gene fragments representing uncultured microbes, or not assigned to known genera in the El Verde soil sequence pools, accounted for 33.1% and 27.3% of the total 16S rRNA gene fragments, respectively. The remaining 16S rRNA gene fragments were assigned to *Acidothermus* (3.1%), ‘*Ca.* Solibacter’ (2.0%) and ‘*Ca.* Udaeobacter’ (7.4%). Taxa with less than 2% representation in the sequence pools are grouped as ‘Others’ and accounted for 27% of the 16S rRNA gene fragments. Following six consecutive transfers (3%, v/v) of the El Verde soil microcosm in defined basal salt medium amended with pyruvate, H_2_, and N_2_O, all 16S rRNA amplicon sequences could be assigned to *Serratia* (68.0%), *Desulfosporosinus* (24.3%), *Desufitobacterium* (7.5%), *Caproiciproducens* (0.19%), *Peptoclostridium* (<0.05%) and *Lachnoclostridium* (<0.05%). Following fifteen sequential transfers in the same medium reduced the microbial diversity to two populations, a *Serratia* sp. and *Desulfosporosinus* sp. In 16S rRNA gene clone libraries generated with DNA collected during Phase II from a 15^th^ generation transfer culture, Sanger sequencing revealed 16S rRNA genes matching those of the *Serratia* sp. and *Desulfosporosinus* sp., and no other sequences were found. Deep metagenome sequencing (total of 387 Gbp obtained) yielded a 2,865-fold coverage of the *Serratia* sp. genome and 15,103-fold coverage of the *Desulfosporosinus* sp. genome (Fig. S3), and 99.48% of the short read sequences could be mapped to the contigs representing these two genomes. Of note, the *Serratia* genomes constructed from co-culture EV and the axenic *Serratia* culture were nearly identical (ANI 99.9%), suggesting the sequencing depth was sufficient to study the genomic characteristics in the co-culture. The assembly of 16S rRNA gene fragments generated eight partial and two complete 16S rRNA genes, which shared between 91.1 to 99.5% sequence identity to the respective *Serratia* sp. and *Desulfosporosinus* sp. 16S rRNA gene sequences determined by Sanger sequencing (Fig. S4). Both genomes possess a single 16S rRNA gene, and the observed differences in 16S rRNA sequences were attributed to errors associated with short read assembly and sequencing. Phase contrast microscopy performed with co-culture EV suspension samples collected during Phase I and Phase II revealed the presence of the reported *Serratia* sp. (45) and the *Desulfosporosinus* sp. cell morphologies (46). Taken together, these observations support that 15 consecutive transfers yielded a consortium (i.e., a co-culture) comprising two bacterial populations, a *Serratia* sp. and a *Desulfosporosinus* sp.

**Supplementary Text S3: Isolation efforts.** Numerous attempts were made to isolate the N_2_O-reducing *Desulfosporosinus* sp. from co-culture EV. Serial 10-fold dilution-to-extinction series in pH 4.5 basal salt liquid medium amended with 2.5 mM pyruvate, 4.16 mM (nominal) N_2_O, and 4.16 mM (nominal) H_2_ recovered N_2_O reduction activity from 10^-6^ dilution vials; however, the *Serratia* sp. was also present. The omission of pyruvate prevented growth of the *Serratia* sp. but without pyruvate, N_2_O consumption did not commence. Plating of serially diluted culture suspension aliquots on solid Tryptic Soy Agar (TSA) medium yielded uniform colonies. Sanger sequencing of PCR-amplified 16S rRNA genes of four isolated colonies from a 10^-6^ dilution plate yielded identical sequences with 100% sequence identity to the *Serratia* sp. Microscopic analysis revealed motile, short rods about 2 µm long representing *Serratia* cells that grew readily in Tryptic Soy Broth medium. Growth also occurred in defined basal salt pH 4.5 medium amended with pyruvate, but the *Serratia* sp. did not consume H_2_ or N_2_O. Following complete N_2_O consumption in co-cultures grown with 2.4 ± 0.01 mM pyruvate, 4.06 ± 1.16 mM (nominal) of H_2_, and 3.96 ± 1.08 mM (nominal) of N_2_O, *Desulfosporosinus* cells outnumbered *Serratia* cells approximately 5-fold. Microscopic observations also documented this population shift and *Desulfosporosinus* cells (motile rods about 6 µm in length) dominated the cell suspensions following N_2_O consumption. Attempts to obtain isolated colonies of the *Desulfosporosinus* sp. in soft agar (0.8% low melting agarose, w/v) shake tubes were not productive because the reduction of N_2_O generated N_2_ gas bubbles preventing the recovery of isolated colonies (Fig. S5). Despite extensive efforts, the *Desulfosporosinus* sp. could not be separated from the *Serratia* sp.

**Supplementary Text S4: Basic genomic features of the populations in co-culture EV.** For sequencing the ‘*Ca.* Desulfosporosinus nitrousreducens’ genome, genomic DNA was isolated from co-culture EV following complete consumption of N_2_O, and the genome constructed from the co-culture metagenome dataset comprising 387 Gbp of raw reads. Construction of the *Serratia* sp. strain MF genome used pure culture DNA and 494 Gbp of raw reads.

Based on CheckM (24) estimates, the completeness of the *Serratia* and *Desulfosporosinus* genomes reached 99.8% and 99.5%, respectively (Table S7). *Serratia* sp. strain MF and *Desulfosporosinus* sp. strain PR have 5,118,938 bp and 5,591,411 bp genomes, G+C contents of 58.8% and 44%, harbor 4,690 and 5,312 protein-coding genes, and possess 81 and 40 tRNA, and 10 and eight rRNA gene sequences, respectively. Genome-wide calculation of the average amino acid identity (AAI) showed that *Serratia* sp. strain MF shares 99.7% AAI with *Serratia marcescens* strain UMH3, suggesting these isolates are closely related. *Desulfosporosinus* sp. strain PR showed highest genomic relatedness with *Desulfosporosinus acidiphilus* strain SJ4 (79.5% AAI) and *Desulfosporosinus acididurans* strain M1 (79.4% AAI). *Desulfosporosinus* species are known for anaerobic sulfate reduction capacity; however, *Desulfosporosinsus* sp. strain PR has an incomplete dissimilatory sulfate reduction pathway, and genes for assimilatory sulfate reduction were absent. Gene clusters encoding two different nitrogenase complexes (*anf* and *nif* gene clusters) are exclusive to the *Desulfosporosinsus* sp. strain PR genome (Fig. S5), indicating the ability to fix N_2_. Albeit the nitrogen fixation capability has never been experimentally validated in members of the genus *Desulfosporosinus*, their genetic makeup (i.e., complete *nif* gene clusters) suggest this to be a shared capability. Also present on the *Desulfosporosinsus* sp. strain PR genome is a *nor* gene cluster encoding nitric oxide (NO) reductase. Genes encoding nitrate and nitrite reductase are absent, consistent with the inability of *Desulfosporosinsus* sp. strain PR to utilize nitrate or nitrite as electron acceptors, although nitrate reduction was reported in some *Desulfosporosinsus* isolates (46-48).

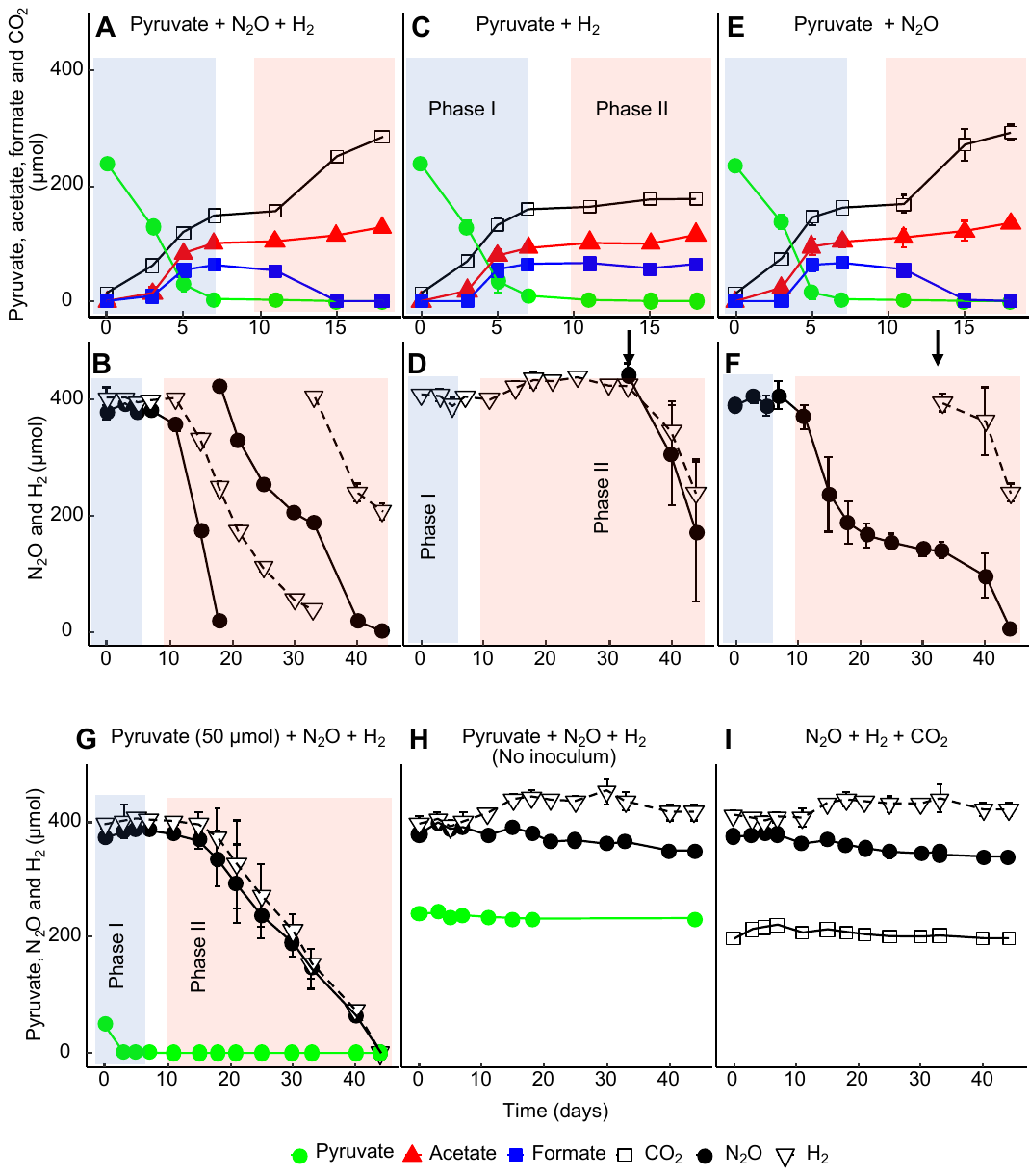
Fig. S1. Phenotypic characterization of co-culture EV. Cultures were grown in 160 mL glass serum bottles containing 100 mL of medium amended with pyruvate (50 or 250 µmol), H_2_ (416 µmol) and N_2_O (416 µmol) (A and B), pyruvate and H_2_ (C and D), or pyruvate and N_2_O (E and F). The arrows in panels D and F indicate additional doses of N_2_O or H_2_ to corresponding culture vessels lack N_2_O (D) or H_2_ (F) to resume N_2_O reduction activity. Pyruvate consumption and associated product formation are documented in panels A, C, and E. N_2_O and H_2_ consumption are documented in panels B, D, F. Panel G shows the performance of co-culture EV grown with a lower amount of pyruvate. Panel H depicts pyruvate, H_2_ and N_2_O amounts over time in vessels without inoculum. Autotrophic activity in cultures with CO_2_, but lacking pyruvate, was not observed (I). The light blue shaded areas represent the pyruvate fermentation phase (Phase I) and the light red areas represent the N_2_O reduction phase (Phase II). The data represent the averages of triplicate incubations and error bars represent the standard deviations (n=3). Error bars are not shown when smaller than the symbol.

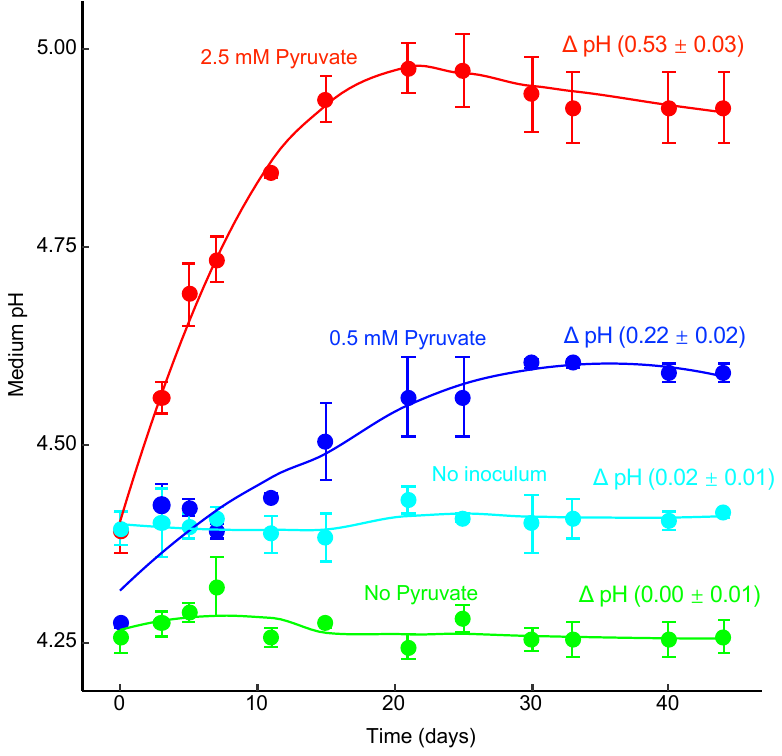

Fig. S2. Medium pH over the course of the incubation. pH profiles were measured in replicate co-culture EV incubation vessels that received 0, 0.5, and 2.5 mM pyruvate. pH changes (Δ pH) were calculated by subtracting the pH values measured immediately following inoculation from pH values measured at the end of incubation period. Pyruvate was not consumed in vessels that were not inoculated with co-culture EV (Fig. S1 H). The lines are plotted based on fitting linear models using the ggplot2 package. The data shown are the averages of triplicate incubations and error bars represent the standard deviations (n=3). Error bars are not shown when smaller than the symbol.

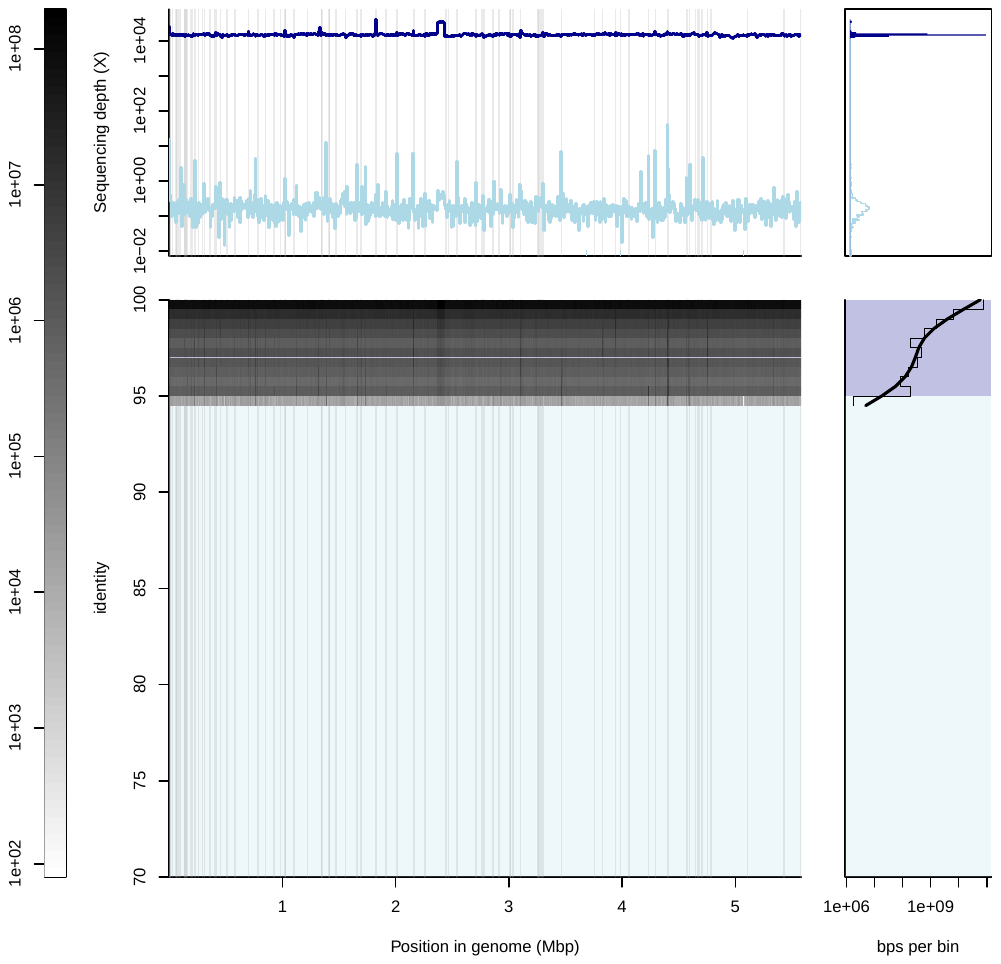

**A**

**C**

**E**

**B**

**D**

**Fig. S3.** Fragment recruitment plot of the ‘*Ca*. Desulfosporosinus nitrousreducens’ strain PR genome to the metagenome dataset derived from the 15^th^ generation co-culture EV. The recruitment was performed via processing a BLAST search of the metagenome fragments against the ‘*Ca*. Desulfosporosinus nitrousreducens’ strain PR genome. The tabular BLAST result was parsed using BLASTab.catsbj.pl, and graphical representation was generated with the BlasTab.recplot2.R embedded in Enveomics collection (30). (A) The bar on the left shows the number of fragments with different identity recruited to each position on the genome. (B) indicates the sequencing depth across the ‘*Ca*. Desulfosporosinus nitrousreducens’ strain PR genome on a logarithmic scale. (C) Sequencing depth histogram with peaks from values above 95% identity automatically identified as skewed normal distribution. (D) Metagenome fragments recruited to the ‘*Ca*. Desulfosporosinus nitrousreducens’ strain PR genome, placed by location (x-axis) and identity (y-axis). (E) Identity histogram of mapping fragments (light gray) and smoothed spline (black). The backgrounds in panels D and E, and the line colors in panels B and C, correspond to identity matches above (dark blue) and below (light blue) 95 %. A recruitment plot of the ‘*Ca*. Desulfosporosinus nitrousreducens’ strain PR genome to the metagenome datasets derived from El Verde soil is not shown as the covered fraction of the 5.6 Mbp genome was below 10% (i.e., 0.56 Mbp).

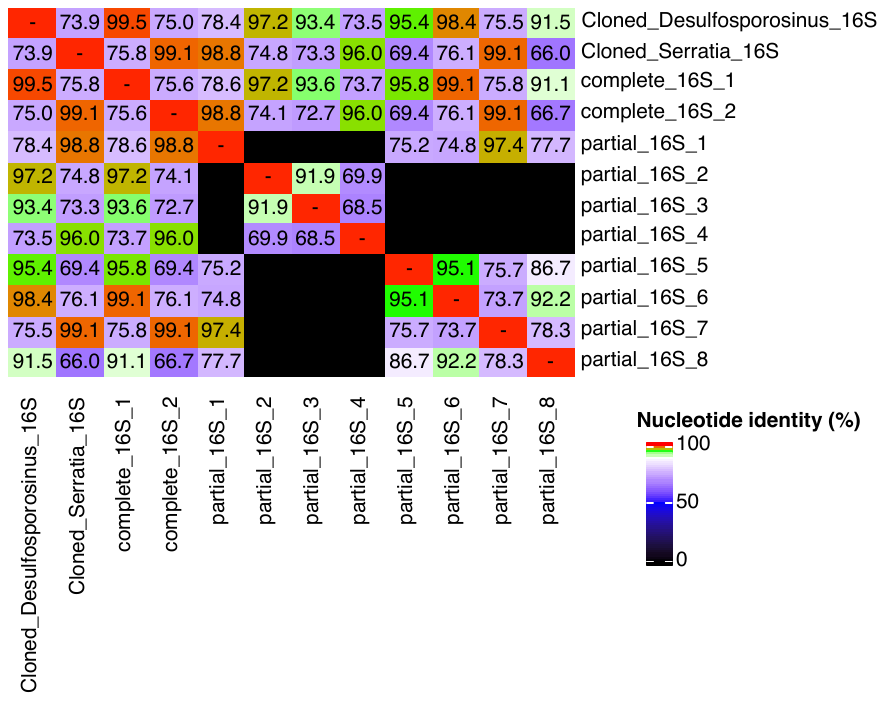

**Fig. S4.** Comparison of ‘*Ca.* Desulfosporosinus nitrousreducens’ strain PR and *Serratia* sp. strain MF 16S rRNA gene sequences derived from cloned 16S rRNA gene fragments (Sanger sequencing) and co-culture EV (metagenome sequencing). DNA extracted from co-culture EV following N_2_O consumption served as template for the amplification of 16S rRNA gene fragments with general primer pair 8F-1541R. 16S rRNA gene fragments were cloned in *E. coli* and two uniform clone populations were obtained, represented by Cloned_Desulfosporosinus_16S and Cloned_Serratia_16S. The assembly of metagenomic reads obtained from a 15^th^ generation co-culture EV yielded full-length 16S rRNA genes of *Serratia* sp. strain MF (1,538 bp) and ‘*Ca*. Desulfosporosinus nitrousreducens’ strain PR (1,467 bp), and eight partial (< 500 bp)16S rRNA genes.

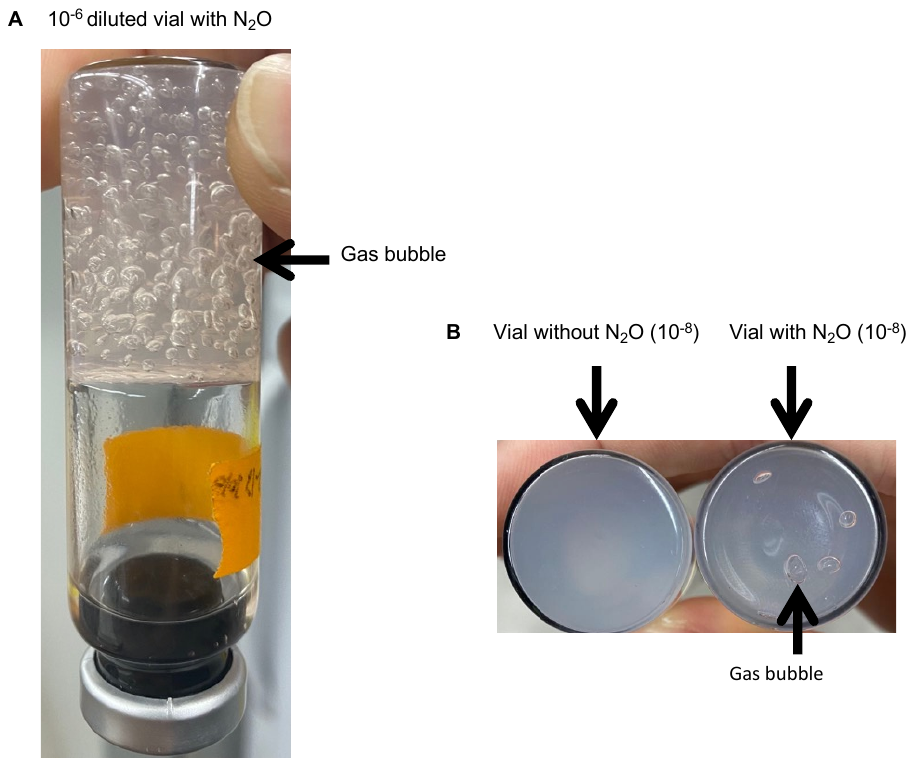

**Fig. S5.** Formation of gas bubbles in soft agar shake tubes. (A) Visible gas bubbles (presumably N_2_) following N_2_O consumption. The vials were incubated with the stoppers down for 28 days. (B) The formation of gas bubbles was strictly dependent on N_2_O, and no bubbles formed in replicate vials without N_2_O. No gas bubbles formed in control incubations without inoculum.

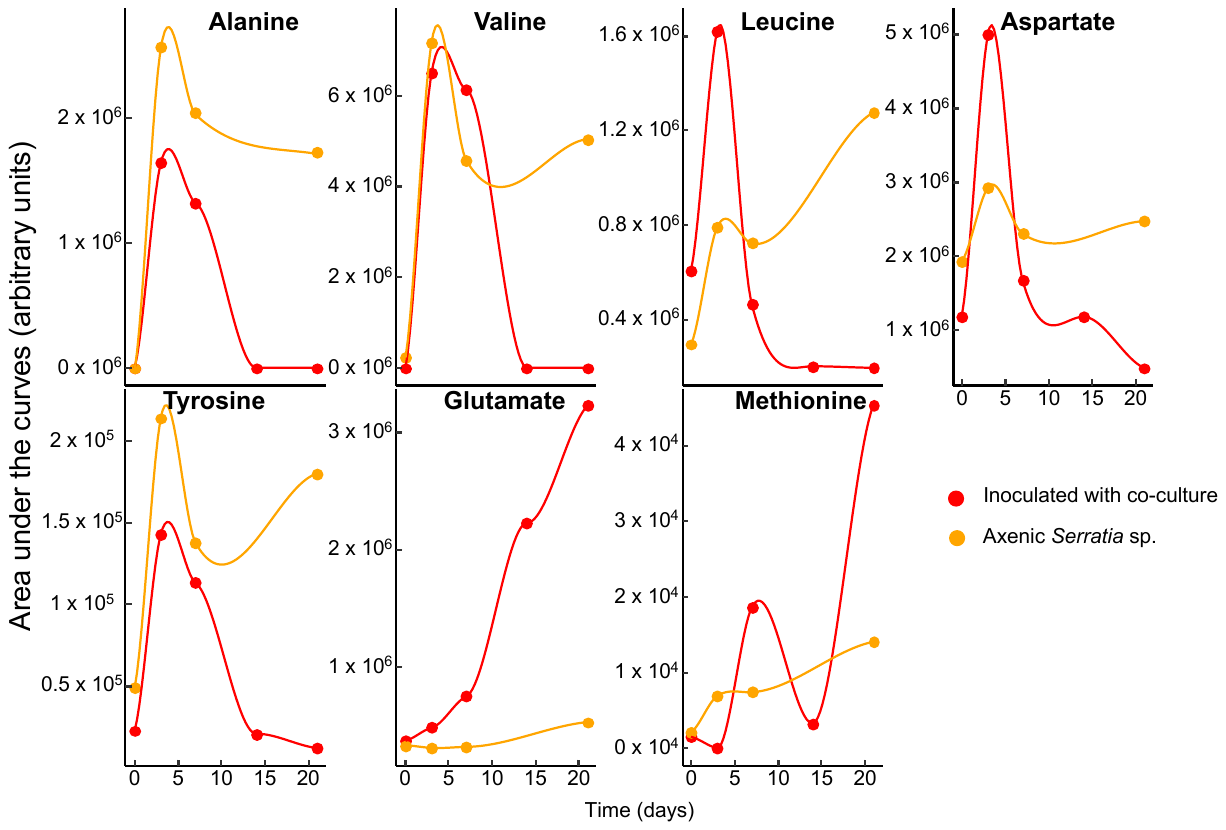

**Fig. S6.** Dynamic changes of amino acids in the supernatants of cultures following inoculation with *Serratia* sp. and with ‘*Ca.* Desulfosporosinus nitrousreducens’ strain PR (as co-culture EV). Axenic *Serratia* sp. cultures amended with pyruvate, N_2_O, and H_2_ were inoculated with co-culture EV on day 7. Orange lines represent amino acids in cultures inoculated with axenic *Serratia* sp. on day 0. Red lines represent amino acids in cultures inoculated with the axenic *Serratia* sp. on day 0 and subsequently inoculated with co-culture EV on day 7. The lines are plotted based on fitting linear models provided in the ggplot2 package. Note that axenic *Serratia* sp. cultures without co-culture EV inoculum cannot reduce N_2_O. Shown are representative data obtained from a single culture in medium reduced with l-cysteine. Cystine was detected in cultures reduced with l-cysteine; however, in samples from replicates from an independent experiment where DTT served as reductant, cystine was not detected, suggesting *Serratia* sp. does not release cystine into the medium.

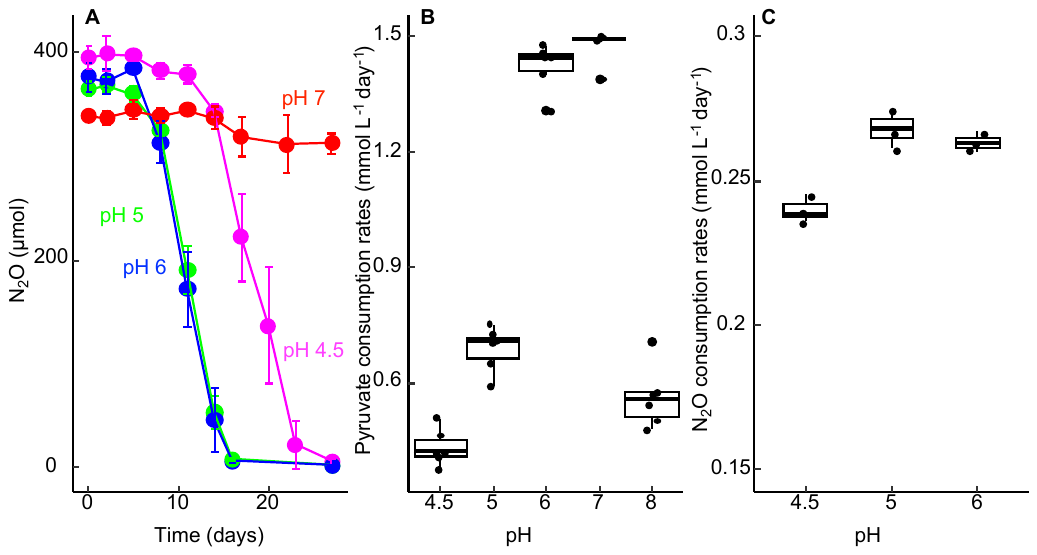

**Fig. S7.** Performance of co-culture EV at different medium pH values. (A) N_2_O reduction was measured in triplicate 160 mL serum bottles containing 100 mL of medium (pH 3.5, 4.5, 5, 6, 7, 8) and inoculated with co-culture EV. N_2_O reduction was observed between pH 4.5 and 6, but not at pH 3.5 and at or above pH 7. (B) Pyruvate consumption rates by *Serratia* sp. in cultures adjusted to pH 4.5, 5, 6, 7, and 8. Pyruvate was consumed in all incubation vessels except for those at pH 3.5. (C) N_2_O consumption rates of ‘*Ca.* Desulfosporosinus nitrousreducens’ at pH 4.5, 5, and 6. No N_2_O consumption was observed in pH 3.5, 7.0, and 8.0 cultures. Consumption rates of pyruvate and N_2_O were calculated using data that fell within the linear ranges of consumption. Each black dot in the box plots represent the consumption rates calculated from a single experiment. The data from three and six replicate cultures were used for the calculations of pyruvate and N_2_O consumption rates, respectively.

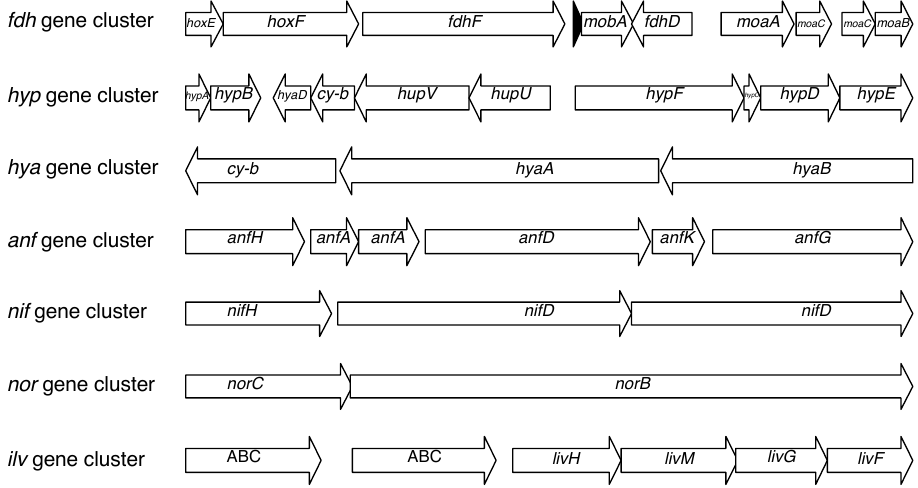

Fig. S8. Gene clusters found on the ‘*Ca*. Desulfosporosinus nitrousreducens’ strain PR genome. Displayed are formate dehydrogenase (*fdh* gene cluster), b-type Ni/Fe hydrogenase (*hyp* gene cluster), group 1 Ni/Fe hydrogenase (*hya* gene cluster), FeFe type Nitrogenase (*anf* gene cluster), Mo-Fe Nitrogenase (*nif* gene cluster), Nitric-oxide reductase (*nor* gene cluster), High-affinity amino acid transport system (*ilv* gene cluster). The arrows indicate operon length and orientations. Preliminary annotation of coding genes was done with Prokka (49), MicrobeAnnotator (50), and the RAST server (29), and genes of interests were curated via blasting to the NCBI nr database. Cy-b: b-type cytochrome; ABC: ATP-binding cassette. The black arrow in the *fdh* gene cluster represents a gene encoding a small protein of unknown function.

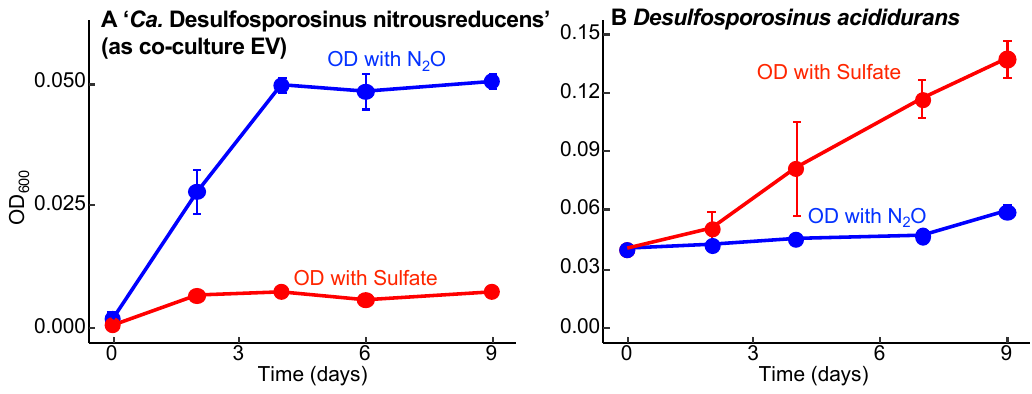

**Fig. S9.** Comparative growth studies with ‘Ca. Desulfosporosinus nitrousreducens’ strain PR (as co-culture EV) and *Desulfosporosinus acididurans* strain D. Panel A shows growth of ‘*Ca*. Desulfosporosinus nitrousreducens’ strain PR with 4.16 mM (nominal) N_2_O (blue) or 5 mM SO_4_^2-^ (red) as electron acceptor in pH 4.5 medium amended with the 15-amino acid mixture and 4.16 mM (nominal) H_2_. Over the 9-day incubation period, strain PR completely consumed the initial dose of N_2_O, whereas cultures amended with 5 mM SO_4_^2-^ showed no growth or SO_4_^2-^ consumption (4.97 ± 0.23 mM sulfate remained). OD values below 0.01 were measured in replicate cultures that did not receive N_2_O as electron acceptor or H_2_ as electron donor. Panel B compares growth of *Desulfosporosinus acididurans* strain D with 4.16 mM (nominal) N_2_O (blue) or 5 mM SO_4_^2-^ (red) as electron acceptors in pH 5.5 medium with 10 mM glycerol as electron donor. Over a 9-day incubation period, *Desulfosporosinus acididurans* strain D reduced 5 mM SO_4_^2-^, but N_2_O was not consumed (4.10 ± 0.59 mM N_2_O remained) in replicate cultures, even after an extended 20-day incubation period. Growth was monitored by measuring the optical density at 600 nm (OD_600_).

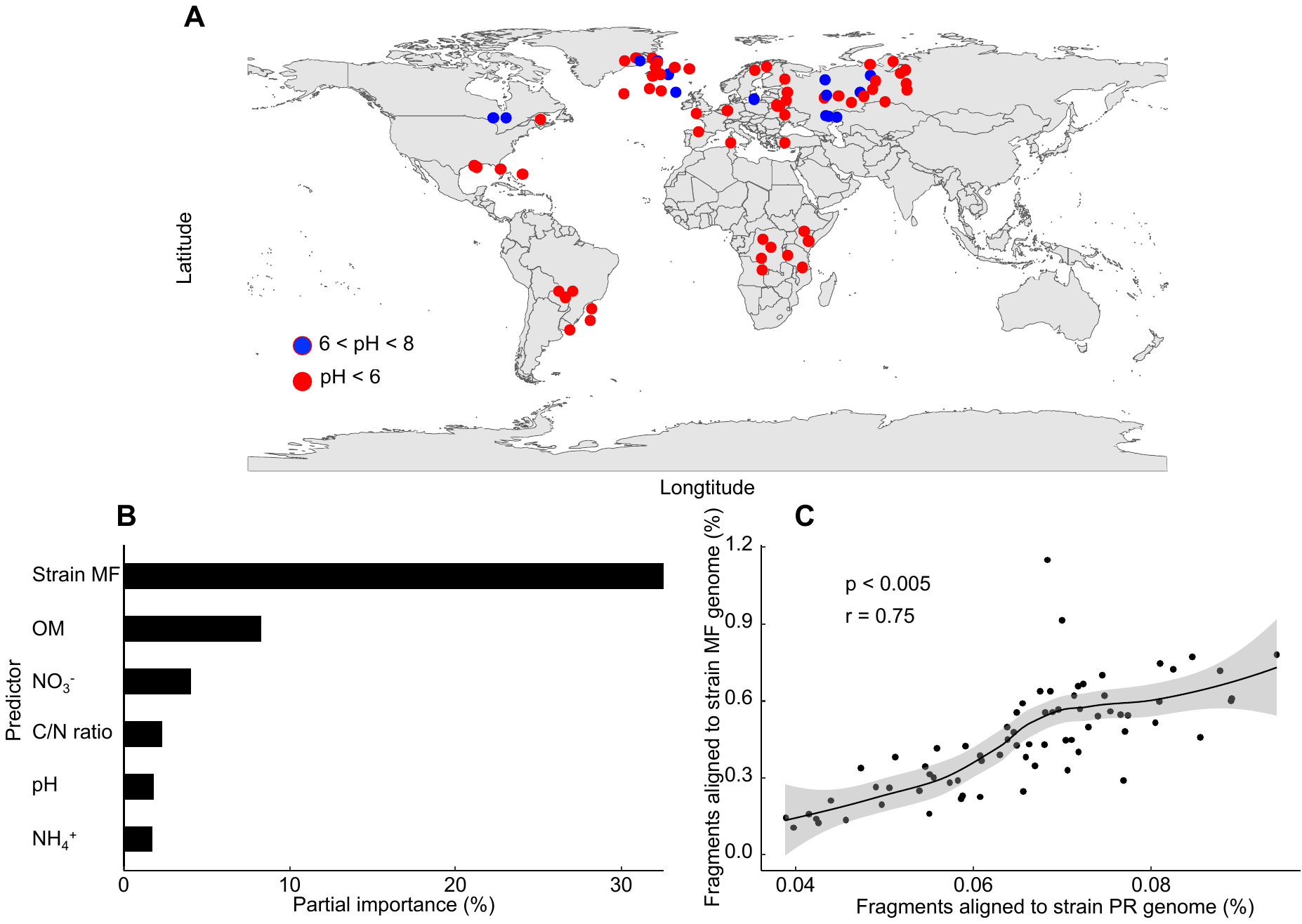

Fig. S10. Distribution of ‘*Ca*. Desulfosporosinus nitrousreducens’ strain PR and *Serratia* sp. strain MF in select publicly available soil metagenomes. Metagenome datasets were downloaded from the European Nucleotide Archive (accession number PRJEB44414) with associated metadata available in a Supplemental Information/Source Data file (https://doi.org/10.1038/s41467-022-29161-3). (A) Geographical distribution of the soil metagenomes included in the analysis. Blue dots indicate a soil pH >7, red dots indicate a soil pH <7. (B) The partial importance (%) of different parameters, including the presence of *Serratia* sp. (based on short reads that align to the *Serratia* sp. strain MF genome), organic matter (OM content), NO_3_^-^ content, NH_4_^+^ content, C/N ratio, and pH for the relative abundance of ‘*Ca*. Desulfosporosinus nitrousreducens’ strain PR based on Random Forest models (51). (C) Correlation between percentages of short read fragments aligned to the *Serratia* sp. strain MF and the ‘*Ca.* Desulfosporosinus nitrousreducens’ strain PR genomes. The significance values for the correlations were calculated with the Pearson test (52) implemented in the ggpubr package (<https://github.com/kassambara/ggpubr>).

**Table S1.** qPCR primers and probe designed for the enumeration of *Serratia* sp. strain MF and ‘*Ca*. Desulfosporosinus nitrousreducens’ strain PR 16S rRNA genes.

| **Target** | **Primer sequence** | **Probe** |
| --- | --- | --- |
| *Serratia* sp. | TGCTACAATTGGCGTATACAA (Ser_1316qF)  GACTACGACATACTTTATGA (Ser_1386qR) | None |
| *Desulfosporosinus* sp. | TGCTACAATGGCCGGTACAG (DS_1316qF)  CGAACTGAGACCGGCTTTCT (DS_1388qR) | FAM-AAGCTGTGAAGTGGAGCCAATC-MGB ^a^ |

^a^ FAM: 6-Carboxyfluorescein attached to the 5’ end of the probe. MGB: Minor Groove Binder that attached to the 3’ end of the probe.

**Table S2.** Sequence information submitted to the NCBI database.

| **Dataset** | **Data type** | **Accession No.** |
| --- | --- | --- |
| Enrichment culture (6^th^ transfer) | Amplicon (16S V3-V4) | SRR24215177 |
| Enrichment culture (9^th^ transfer) | Amplicon (16S V3-V4) | SRR24083098 |
| Co-culture EV (15^th^ transfer) | SRA ^a^ | SRR24709127 |
| *Serratia* sp. (isolate) | SRA ^a^ | SRR24709126 |
| 16S rRNA gene (*Desulfosporosinus*) | Complete gene | OR076434 |
| 16S rRNA gene (*Serratia*) | Complete gene | OR076433 |
| ‘*Ca.* Desulfosporosinus nitrousreducens strain PR’ | GenBank Assembly ^b^ | GCA_030954495.1 |
| *Serratia* sp. strain MF | GenBank Assembly ^b^ | GCA_030954505.1 |

^a^ SRA: Sequence Read Archive (raw sequence data)

^b^ Genome assembly constructed from SRA data

**Table S3.** Organismal and genomic information of microorganisms with reported N_2_O reduction capacity under different pH conditions.

| **Organism** | **Genome NCBI  acc no.** | **Reported pH range for N_2_O reduction** | ***nosZ* clade** | **Reference** |
| --- | --- | --- | --- | --- |
| ‘*Ca*. Desulfosporosinus nitrousreducens’ strain PR | GCA_030954495.1 | 4.5-6 | II | This work |
| *Desulfosporosinus meridiei* strain DSM 13257 | NC_018515.1 | ND | II | (53) |
| *Nitratiruptor labii* strain HRV44 | AP022826.1 | 5.4-6.4 | II | (54) |
| *Anaeromyxobacter dehalogenans* strain 2CP-1 | NC_011891.1 | ~7 | II | (53) |
| *Ferroglobus placidus* | NC_013849.1 | ND | II | (53) |
| *Azospira oryzae* | GCA_905120365.1 | ND | II | (53) |
| *Paracoccus denitrificans* strain DSM 413 | AL591688.1 | 7-8 | I | (55) |
| *Shewanella loihica* strain PV-4 | NC_009092.1 | 6-8 | I | (56) |
| *Alicycliphilus denitrificans* strain I51 | AP024172.1 | 6-9 | I | (57) |
| *Bradyrhizobium diazoefficiens* strain 110*spc*4 | NZ_CP032617.1 | 6-8 | I | (58) |

**Table S4.** Genomes used in phylogenomic analysis and classification of ‘*Ca*. Desulfosporosinus nitrousreducens’.

| **Organism** | **NCBI accession No. of genomes** |
| --- | --- |
| ‘*Ca*. Desulfosporosinus nitrousreducens’ strain PR | GCA_030954495.1 |
| *Desulfosporosinus youngiae* DSM 17734 | NZ_CM001441.1 |
| *Desulfosporosinus orientis* DSM 765 | NC_016584.1 |
| *Desulfosporosinus metallidurans* strain OL | NZ_MLBF01000001.1 |
| *Desulfosporosinus meridiei* DSM 13257 | NC_018515.1 |
| *Desulfosporosinus lacus* DSM 15449 | NZ_FQXJ01000052.1 |
| *Desulfosporosinus hippie* DSM 8344 | NZ_fncp01000067.1 |
| *Desulfosporosinus fructosivorans* strain 63.6F | NZ_SPQQ01000010.1 |
| *Desulfosporosinus acidiphilus* SJ4 | NC_018068.1 |
| *Desulfosporosinus acididurans* strain M1 | NZ_LDZY01000001.1 |
| ‘*Ca.* Desulfosporosinus infrequens’ | NZ_OMOF01000971.1 |
| *Syntrophobotulus glycolicus* DSM 8271 | NC_015172.1 |
| *Desulfitobacterium* *metallireducens* DSM 15288 | NZ_CP007032.1 |
| *Desulfitobacterium* *hafniense* DCB-2 | NC_011830.1 |
| *Desulfitobacterium* *dichloroeliminans* LMG P-21439 | NC_019903.1 |
| *Desulfitobacterium* *dehalogenans* ATCC 51507 | NC_018017.1 |
| *Desulfitobacterium* *chlororespirans* DSM 11544 | NZ_FRDN01000033.1 |
| *Dehalobacter* *restrictus* strain E1 | CANE01000102.1 |
| *Dehalobacter* *restrictus* strain 12DCA | NZ_CP046996.1 |
| *Dehalobacter restrictus* DSM 9455 | NZ_CP007033.1 |

**Table S6.** Amino acids and concentrations used to augment the growth medium for cultivation of ‘*Ca*. Desulfosporosinus nitrousreducens’. ^a^

| **Amino acid** | **Amino acid  concentration (µM)** | **Carbon (µmol 100 mL medium^-1^)** |
| --- | --- | --- |
| Alanine | 56 | 16.8 |
| Valine | 77 | 38.5 |
| Aspartate | 82 | 33.1 |
| Isoleucine | 38 | 22.9 |
| Methionine | 27 | 13.4 |
| Tyrosine | 16 | 14.9 |
| Histidine | 19 | 11.6 |
| Tryptophan | 10 | 10.8 |
| Glutamate | 68 | 34.0 |
| Proline | 86 | 43.5 |
| Arginine | 28 | 17.2 |
| Glycine | 40 | 8.0 |
| Threonine | 42 | 16.8 |
| Lysine | 69 | 41.1 |
| Serine | 76 | 22.9 |
|  |  | Total carbon 383.7 |

^a^ The addition of individual amino acids did not stimulate N_2_O-dependent growth of ‘*Ca*. Desulfosporosinus nitrousreducens’. Similarly, a 5-amino acid mixture comprising alanine, valine, leucine, aspartate, and tyrosine did not promote N_2_O consumption. Weak and incomplete N_2_O reduction was observed in cultures that received a 6-amino acid mixture comprising alanine, valine, leucine, aspartate, tyrosine, and methionine. The 15-amino acid mixture (i.e., all amino acids listed in Table S3) supported growth of ‘*Ca*. Desulfosporosinus nitrousreducens’ via hydrogenotrophic N_2_O reduction.

**Table S7.** Genome features of ‘*Ca.* Desulfosporosinus nitrousreducens’ strain PR and *Serratia* sp. strain MF, and comparison to closest relatives.

|  | **Organism ^a^** | | | | |
| --- | --- | --- | --- | --- | --- |
| **Feature** | **(1)** | **(2)** | **(3)** | **(4)** | **(5)** |
| Genome size (bp) | 5,591,411 | 4,991,181 | 4,637,866 | 5,118,938 | 5,300,955 |
| Completeness (%) ^b^ | 99.5 | 99.1 | 92.3 | 98.0 | 95.6 |
| Contamination (%) ^b^ | 5.7 | 0.0 | 5.14 | 0.53 | 4.03 |
| GC content (%) | 44 | 42 | 41.5 | 58.8 | 59.5 |
| 5S rRNA genes | 8 | 8 | 8 | 8 | 8 |
| 16S rRNA genes | 1 | 9 | 10 ^d^ | 1 | 7 |
| 23S rRNA genes | 1 | 8 | 11 ^d^ | 1 | 7 |
| tRNA genes | 105 | 66 | 67 | 81 | 91 |
| Coding sequences | 5,312 | 4,554 | 4,317 | 4,690 | 4,941 |
| Accession number | GCA_030954495.1 | GCA_000255115.3 | GCA_001029285.1 | GCA_030954505.1 | GCF_002220655.1 |

^a^ (1) ‘Ca. Desulfosporosinus nitrousreducens’ strain PR (this study)

(2) *Desulfosporosinus acidiphilus* strain SJ4 (59)

(3) *Desulfosporosinus acididurans* strain M1 (46)

(4) *Serratia* sp. strain MF (this study)

(5) *Serratia marcescens* strain UMH3 (60)

^b^ Genome completeness and contamination were evaluated based on the presence of 120 single copy genes using CheckM.

^c^ Only partial 16S and 23S rRNA genes were found on *Desulfosporosinus acididurans* strain M1 genome.
